# Origin and modern microbial ecology of secondary mineral deposits in Lehman Caves, Great Basin National Park, NV, USA

**DOI:** 10.1101/2023.08.15.553329

**Authors:** Zoë E. Havlena, Louise D. Hose, Harvey R. DuChene, Gretchen M. Baker, Doug Powell, Amanda L. Labrado, Benjamin Brunner, Daniel S. Jones

## Abstract

Lehman Caves is an extensively decorated, high desert cave that represents one of the main tourist attractions in Great Basin National Park, Nevada. Although traditionally considered a water table cave, recent studies identified abundant speleogenetic features consistent with a hypogenic and, potentially, sulfuric acid origin. Here, we characterized white mineral deposits in the Gypsum Annex (GA) passage to determine whether these secondary deposits represent biogenic minerals formed during sulfuric acid corrosion, and explored microbial communities associated with these and other mineral deposits throughout the cave. Powder X-ray diffraction (pXRD), scanning electron microscopy with electron dispersive spectroscopy (SEM-EDS), and electron microprobe analyses (EPMA) showed that, while most white mineral deposits from the GA contain gypsum, they also contain abundant calcite, silica, and other phases. Gypsum and carbonate-associated sulfate isotopic values of these deposits are variable, with δ^34^S_V-CDT_ between +9.7‰ and +26.1‰, and do not reflect depleted values typically associated with replacement gypsum formed during sulfuric acid speleogenesis. Petrographic observations show that the sulfates likely co-precipitated with carbonate and SiO_2_ phases. Taken together, these data suggest that the deposits resulted from later stage meteoric events and not during an initial episode of sulfuric acid speleogenesis. Most sedimentary and mineral deposits in Lehman Caves have very low microbial biomass, with the exception of select areas along the main tour route that have been impacted by tourist traffic. High-throughput 16S rRNA gene amplicon sequencing showed that microbial communities in GA sediments are distinct from those in other parts of the cave. The microbial communities that inhabit these oligotrophic secondary mineral deposits include OTUs related to known ammonia-oxidizing *Nitrosococcales* and Thaumarchaeota, as well as common soil taxa such as Acidobacteriota and Proteobacteria. This study reveals microbial and mineralogical diversity in a previously understudied cave and expands our understanding of the geomicrobiology of arid, hypogene cave systems.

## 1. INTRODUCTION

Earth’s subsurface may contain as much as 70% of total cells on the planet (Magnabosco et al., 2018; Magnabosco et al., 2019), and the microorganisms that inhabit this expansive and diverse biosphere are important for rock weathering, sedimentary carbon cycling, and many other critical biogeochemical processes (e.g., Engel & Randall, 2011; Puente-Sánchez et al., 2014; Sabuda et al., 2020). However, most of the subsurface is inaccessible to human exploration, and as a result, we know much less about these deep microbial communities than we do about those at the surface, especially in the continental subsurface.

We can directly access subterranean microbial activity through caves. In caves, microorganisms facilitate the cycling of elements like sulfur, nitrogen, and iron, and chemolithoautotrophs can drive primary productivity in these sometimes nutrient-limited ecosystems (Jones & Macalady, 2016). Cave microorganisms cycle carbon and can control fluxes of greenhouse gases (Webster et al., 2018; Martin-Pozas et al., 2022), and microbial activity can directly or indirectly affect speleothem formation and precipitate secondary minerals (Northup & Lavoie, 2001). The study of these subsurface microbiological processes in caves not only enhances our understanding of biogeochemical cycles on Earth but also provides insights into the potential for life in similar environments on other planets (Boston et al., 2001).

Lehman Caves is a popular show cave in Great Basin National Park, Nevada, USA, that is visited by over 33,000 visitors a year (Great Basin National Park, 2019). The cave has also been a valuable source of speleothem records on the paleohydrology and paleoclimate of the Southwest (Shakun et al., 2011; Lachniet et al., 2014; Cross et al., 2015; Steponaitis et al., 2015; Lachniet et al., 2016). However, the geology and origin of the cave itself has not been described in detail. While historically considered an epigenic cave that formed at the water table (Moore & Sullivan, 1978), this explanation is not consistent with the arid setting of Lehman Caves and lack of connection to ancient or modern drainage features. Recently, Hose et al. (2021) reinterpreted features within Lehman Caves as evidence for hypogene, sulfuric acid speleogenesis, contrasting the classic interpretation with a complex, multistage formational history.

Hypogenic caves form by “bottom-up” processes, where carbonate dissolution is driven by rising groundwaters charged with carbon dioxide (CO_2_), hydrogen sulfide (H_2_S), or other acids. In “sulfidic” hypogenic caves, voids form where H_2_S-rich groundwaters are exposed to oxygen, usually at or near the water table. Where O_2_ and H_2_S interact, H_2_S oxidizes to sulfuric acid (H_2_SO_4_), which dissolves the carbonate bedrock. This process is known as “sulfuric acid speleogenesis” (SAS) and is responsible for large and spectacular cave systems, including Carlsbad Cavern and Lechuguilla Cave in New Mexico and the Frasassi Caves in Italy (DuChene et al., 2017; Galdenzi & Maruoka, 2003). In Lehman Caves, Hose et al. (2021) proposed that passages were first developed and enlarged in an initial SAS phase, and subsequently modified during a later stage of meteoric water infiltration, where extensive carbonate speleothems overprinted most early speleogenetic features. SAS morphologies in the cave include acid pool features, bubble trails, hollow coralloidal stalagmites, and irregularly shaped pseudoscallops (Hose et al., 2021). These features are most evident in a passage known as the Gypsum Annex (GA), where extensive, loose, white mineral residues distinguish it from the other highly decorated passages, perhaps because the GA was protected from infiltration by an overlying quartzite caprock (Hose et al., 2021).

During sulfuric acid speleogenesis, gypsum (CaSO_4_•2H_2_O) crusts form on cave walls where H_2_S degassing and oxidation drives sulfuric acid corrosion subaerially (Egemeier, 1981). As this process continues, microcrystalline gypsum precipitates on the cave walls and ceilings, and these secondary mineral crusts eventually detach and accumulate as “breakdown” on the cave floor (Galdenzi & Maruoka, 2003; Hose et al., 2000). If gypsum deposits are protected from dissolution, they can persist for hundreds of thousands to millions of years in ancient sulfuric acid caves (Hill, 1990; Polyak et al., 1998; Jagnow et al., 2000; Galdenzi & Jones, 2017).

Gypsum also occurs in other caves. Most limestone caves form by “top-down” CO_2_-driven dissolution processes (epigenic karst) (Palmer, 1991). Secondary gypsum can form in epigene caves, often by precipitation from high sulfate seepage or other cave waters, where sulfate is sourced from nearby evaporites or pyrite-containing bedrock (e.g., Palmer & Palmer, 2003; Metzger et al., 2015). While these gypsum deposits are typically not as extensive as those formed during sulfuric acid speleogenesis, diagnosing their origin can be challenging. Stable sulfur isotope ratios may provide additional information. δ^34^S values of SAS gypsum reflect the dissolved H_2_S, and are very light when the H_2_S results from sulfate reduction. For example, gypsum deposits in the Frasassi Caves have δ^34^S between −7‰ and −20‰ (Galdenzi & Maruoka, 2003), and even lighter gypsum of −23‰ to nearly −30‰ has been reported for other SAS origin caves (Cueva de Villa Luz and Guadalupe Mountain caves, respectively; Hose et al., 2000; Hill, 2000).

In active sulfuric acid caves, gypsum formation is driven by biological activity. Chemolithotrophic sulfide-oxidizing bacteria and archaea catalyze H_2_S oxidation much faster than abiotic rates (Engel et al., 2004; Mansor et al., 2018), speeding up sulfuric acid formation and inducing gypsum replacement. Microorganisms contribute to the formation of other secondary minerals in both hypogene and epigene caves, such as elemental sulfur, iron and manganese-rich crusts, or carbonate moonmilk (Northup & Lavoie, 2001; Spilde et al., 2005, Cañaveras et al., 2006, Jones et al., 2012, Carmichael et al., 2013; Engel et al., 2013, Hamilton et al., 2015). Slow-growing microbial communities that occur in oligotrophic cave environments may still contribute to bedrock corrosion and secondary mineral formation throughout the long lifetime of caves (Northup & Lavoie, 2001; Barton & Northup, 2007). However, we don’t know if microbial associations with gypsum persist in ancient SAS systems, or whether and which microbes colonize gypsum deposits in epigenic caves.

The white mineral deposits on the walls and floors of the GA passage resemble speleogenetic gypsum crusts common to SAS caves. However, they could also be later-stage deposits that formed after an initial SAS episode. Therefore, the first goal of this study was to explore the mineralogy and isotopic composition of deposits from the GA and other areas of Lehman Caves, to evaluate the origin of these secondary minerals. The second goal was to describe modern microbial colonization of mineral surfaces throughout the cave. Unlike the energy-rich environment of active SAS caves, relict karst systems often have limited energy resources and sparse microbiota (Barton & Jurado, 2007). However, slow microbial processes may still contribute to bedrock corrosion and secondary mineral formation throughout the lifetime of the cave. Whether, and how, microbial communities continue to use gypsum and other secondary precipitates has implications for the microbial ecology and biosignature preservation in other terrestrial sulfates, or in geochemically analogous deposits observed on the surface of Mars (Hays et al., 2017). To our knowledge, this is the first detailed study on the geochemistry of mineral deposits in the GA, and the first comprehensive geobiology survey of Lehman Caves.

## 2. MATERIALS AND METHODS

### 2.1 Study area and sample collection

Lehman Caves (39.0054° N, 114.2207° W) is in Great Basin National Park in east-central Nevada (Figure 1A). Despite the name being plural, Lehman Caves is one connected cave system, and the name does not refer to any of the other caves within the boundaries of Great Basin National Park (Prudic et al., 2015). The cave was designated as a National Monument in 1922, and was incorporated into Great Basin National Park in 1986. Tourist traffic occurs along a 1-km ranger-guided trail that currently only goes as far as Grand Palace and Sunken Garden. From 1961-1981, tours of the main cave looped through the Talus Room and Royal Gorge as well. Lehman Caves is the longest explored cave in Nevada, and contains approximately 3 km of horizontal passage in the locally metamorphosed mid-Cambrian Pole Canyon Limestone (PCL). The PCL was metamorphosed to a mylonitic marble during the Miocene Snake Range décollement event, prior to the development of the cave (Hose et al., 2021). Small outcrops of the mid-Cambrian Pioche Shale are observable where it intruded into the PCL during a period of local faulting, and part of the cave is capped by the Neoproterozoic Prospect Mountain Quartzite (PMC) (Figure 1C). While the age of the cave is not currently well constrained, it is probably many million years old. Disequilibrium uranium-series dating shows that one Lehman speleothem is older than 2.2 Ma, the limit of ^234^U/^238^U disequilibrium methods (Great Basin National Park, 2019), and it is likely much older than that. For a more detailed site description, regional geologic context, and discussion of possible age, readers are referred to Hose et al. (2021).

**Figure 1.**
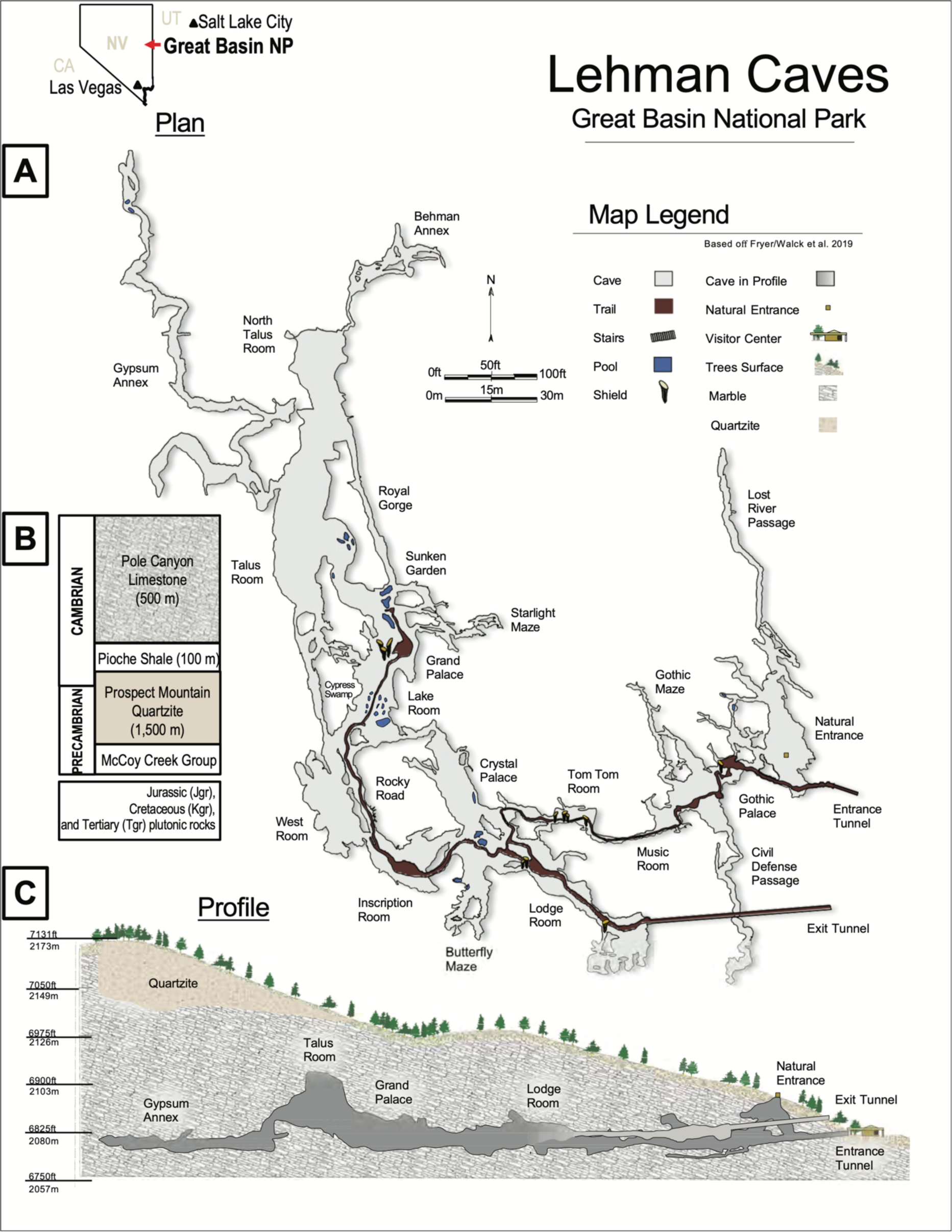
(A) Annotated map of Lehman Caves, by Shane Fryer, Cynthia Walck, and others (*unpub.,* Great Basin National Park map archive). (B) Relevant stratigraphic units and (C) profile of Lehman Caves, adapted by Hose et al. (2021), showing where an outcrop of Prospect Mountain Quartzite (PMQ, in tan) was mapped above the Gypsum Annex.

Most samples for this study were collected from the Gypsum Annex (GA) passage (Figure 1). The GA was discovered in 1952 (Trexler, 1966), and branches off from the Talus Room, where the passage is accessible by a short cable ladder. The GA has never been part of the cave open to tour or for recreational use. To our knowledge, the only prior mineralogy study of the GA is limited to a brief unpublished report to Great Basin National Park of observations by K.J. Murata of the United States Geological Survey (Anonymous, 1965). As described earlier, the GA preserves features recently reinterpreted as having formed under hypogenic, sulfuric acid speleogenesis (Hose et al., 2021). There is no detectable H_2_S currently present in air or waters in the GA or any other part of the cave. Sulfide gases were noted by smell during extensive sampling of surficial waters in the Lehman Creek drainage by the United States Geological Survey (USGS) in 2004 (Cook & Corboy, 2004), but this was attributed to reduction of sulfate ions in the springs and it is unlikely to reflect any modern sulfide-bearing groundwaters.

Samples were collected in 15- or 50-ml sterile tubes using aseptic techniques during trips in September 2018 and June 2019. Samples are summarized in Table 1. Sampling was done conservatively so as to minimize disturbance. Samples were subdivided into fractions for mineralogical analysis, DNA analysis, and enrichment culturing. Samples for DNA extraction were immediately frozen on dry ice in the field and subsequently stored at −80 °C upon return to New Mexico Tech, samples for culturing were stored at 4 °C, and samples for microscopy were fixed in 4% PFA as in Jones et al. (2017a) and stored at 4 °C. In 2019, eight water samples were collected from pools in GA, Talus Room, and near the main tour trail. Water samples were syringe filtered with 0.2 μm PTFE filters (VWR, Radnor, PA), and preserved with either 6N hydrochloric acid (HCl) or 5% w/v zinc acetate for major ion and dissolved sulfide analyses, respectively. Conductivity, temperature, and pH were measured on site using an Oakton PCSTestr 35 field probe.

**Table 1.**
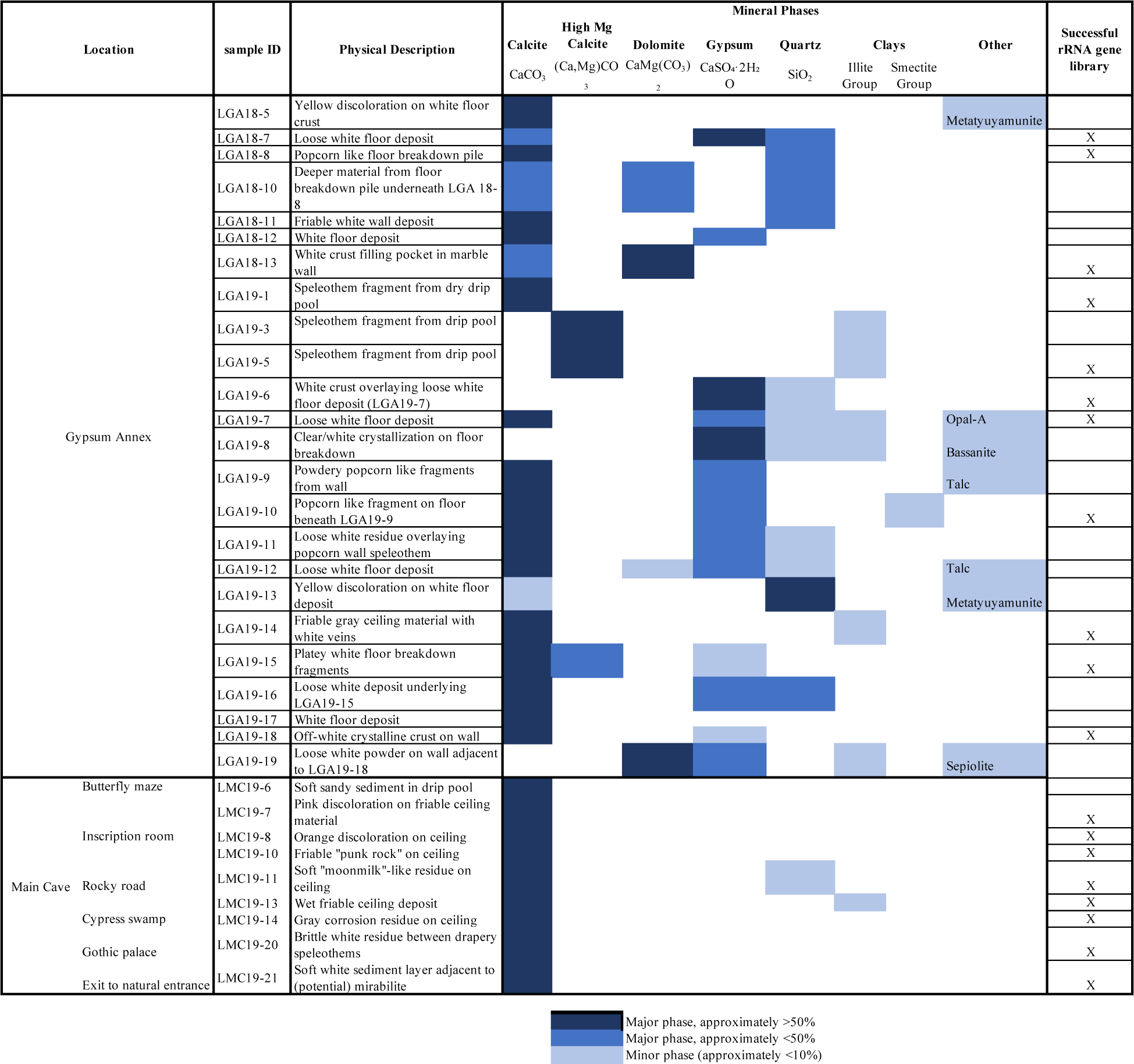
Sample description and mineralogy.

### 2.2 Geochemical and Mineralogical Analyses

Sediment samples for mineralogical and geochemical analyses were oven dried at 60 °C for 36 hours. A small portion (0.5-1g) of the dried material was ground using a ceramic mortar and pestle and analyzed by powder X-Ray Diffraction (pXRD) with a Panalytical X’Pert Pro Diffractometer with a Cu Kα source from 5° to 70° (2θ) for 300 s at 45 kV and 40 mA. The phases in resulting diffractograms were examined and relative proportions of major elements were obtained and visualized with the Panalytical High Score Plus phase analysis software using International Centre for Diffraction Data (ICDD) reference patterns for calcite (01-085-1108), gypsum (01-070-0983), and quartz (01-070-0983).

Some samples were imaged uncoated with a Hitachi TM-1000 tabletop scanning electron microscope (Hitachi, Tokyo, Japan) equipped with an energy-dispersive spectrometer (EDS). Two additional samples with yellow coloration were sputter coated with platinum and analyzed by scanning electron microscopy and energy dispersive X-ray spectroscopy (SEM-EDS) using a Field Emission JEOL IT-700HR. Analysis was done using an Oxford X-maxN EDS and Oxford AzTec software. Thin sections were prepared from two intact fragments from GA white floor deposits (Wagner Petrographic, Lindon, UT). Petrographic analysis of thin sections was performed with Olympus CX31 and BX53 petrographic microscopes. Electron microprobe analysis (EMPA) of thin sections was performed with a Cameca SX-100 electron microprobe with 4 WDS spectrometers using an acceleration voltage of 15 kV, beam current of 10 nA, and a 5 or 20 µm spot size for quantitative analyses. WDS spectra and X-Ray maps were performed using a 20nA beam current with a point beam. X-ray mapping was performed for Si, Mg, Ca, and S. All data was collected and processed using Cameca PeakSight 6.2.

Dissolved elements in water samples, including major cations, total P, and total S, were determined by inductively coupled plasma optical emission spectroscopy (ICP-OES) on an Agilent 5100 ICP-OES. Mineral saturation indices were calculated using PHREEQC version 3 in R (Parkhurst & Appelo, 2013; Charlton et al., 2023) with the PHREEQC database, assuming that all dissolved sulfur was present as sulfate, and using Cl^−^ for charge balance.

### 2.3 Stable isotope analyses

We measured the stable sulfur isotope composition (δ^34^S) of sulfate in gypsum and carbonate-associated sulfate (CAS) in ten samples from the GA and two samples from other sections of the cave. Samples were prepared following the methods of Gischler et al. (2020). Samples were ground to a fine powder using a ceramic mortar and pestle. The NaCl-soluble easily soluble sulfate (ESS) fraction was obtained from 0.5 to 3.0 g of powdered sample in pre-weighed 50 ml Falcon tubes, which was dissolved by shaking horizontally in 35 ml of 10% NaCl for 48 h. The resulting supernatant was separated by centrifugation, and dissolved sulfate within the supernatant was precipitated by adding 200 μL 6N HCl, incubating for 3 minutes, and then adding 5 mL of 1M barium chloride (BaCl_2_) and shaking overnight. The BaSO_4_ precipitate was collected by centrifugation and separated into a pre-weighed 2 ml microfuge tube. The pellet was then washed with deionized water until pH was >5, and the pellet was dried at 60 °C for 24 hours. For CAS, the HCl soluble fraction was obtained from the residual pellet after NaCl dissolution. 5N HCl was added to the pellet in 100 μL increments, until the sample stopped reacting (usually after at least 2 mL were added). Deionized water was added to bring the volume to 35 ml, and the sample was centrifuged and supernatant was filtered (0.45 μm, PES) into a new 50 ml tube. The sulfate liberated from the carbonate during acidification was precipitated with BaCl_2_ as described above.

δ^34^S isotopic analysis was performed by isotope ratio mass spectroscopy at The University of Texas at El Paso using a Pyro Cube Elemental Analyzer and GeoVisION isotope ratio mass spectrometer (EA-IRMS). Isotope ratios for ESS (δ^34^S_ESS_) and CAS (δ^34^S_CAS_) are reported using standard delta notation relative to the Vienna Canyon Diablo Troilite (V-CDT) reference material,

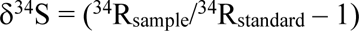

where ^34^R_sample_ = ^34^S/^32^S, where δ^34^S values are reported in ‰.

### 2.4 DNA extraction and rRNA gene library preparation

We compared different methods for extracting DNA from the gypsum-rich samples. After initial challenges extracting DNA from Lehman Caves samples collected in 2018, we evaluated different commercial DNA extraction kits and procedural modifications by extracting Lehman Caves samples as well as “spike in” experiments in which cells from either an *Acidithiobacillus thiooxidans* culture or a microbial community standard (ZymoBIOMICS Microbial Community Standard, Zymo Research Corp.) were added to low biomass gypsum collected from the Frasassi Caves.

First, total DNA extractions were performed on select Lehman Caves samples using the DNeasy Powersoil Pro DNA Isolation Kit (Qiagen, Germantown, MD, USA), DNeasy PowerMax Soil Kit, DNeasy PowerBiofilm Kit, and the now discontinued DNeasy Powersoil DNA Isolation Kit (all Qiagen). For extractions with the DNeasy Powersoil Pro kit, the bead-beating step was either performed with a Vortex-Genie 2 (Scientific Industries, Inc., Bohemia, NY, USA) for 5, 10, and 15 min at maximum speed, or using a bead beater (Biospec Products Mini Beadbeater) for 20, 40, and 60 seconds at 2500 RPM. These same kits were then used to extract low biomass gypsum spiked with *Acidithiobacillus* cultures. For these experiments, either 1 ml, 100 µl, or 10 µl of an isolate culture was added to ∼0.25 g low biomass gypsum shavings in 2 ml microfuge tubes, and extracted alongside gypsum only or isolate only controls. Samples were either extracted directly, or first treated with a pre-extraction process in which gypsum was dissolved by horizontally shaking the sample in 1× filter sterilized phosphate buffered saline (PBS) for 60 minutes. The PBS was acidified to pH 2 with HCl for the *Acidithiobacillus* spike-in experiment. In all cases, blanks consisting of kit reagents only were processed in tandem with extractions (“DNA blank”). Efficacy of extraction and amplification was assessed by PCR targeting the V4 hypervariable region of the 16S rRNA gene using primers 515f modified (515f-m; 5’-GTG YCA GCM GCC GCG GTA A-3’) and 806r (5’-GGA CTA CNV GGG TWT CTA AT-3’) (Walters et al., 2016). All PCRs included a negative control PCR reagent blank (“PCR blank”), as well as the DNA blank and a positive control of DNA from an isolate culture.

Following indications that the highest DNA yield was achieved using the PowerSoil Pro kit without prior gypsum removal with PBS, we then tested the fidelity of this kit by extracting DNA from gypsum spiked with cells from a mock community (ZymoBIOMICS Microbial Community Standard, Zymo Research, Irvine, CA, USA). We added 25 μL of this standard (∼4.7×10^8^ cells) to 0.25 g of gypsum, and extracted half of each aliquot using the Vortex-Genie 2 and other half with the bead beater, as described above. All extractions of spiked gypsum were performed in triplicate, alongside aliquots of gypsum only, mock community only, and extraction blanks. Ultimately, the PowerSoil Pro kit with bead beating in intervals was selected and used to extract all samples, including those that initially failed to strongly amplify.

rRNA gene amplicon libraries were prepared following the “in house” methods of Jones et al. (2017a) that are appropriate for low biomass samples. The V4 region of the 16S rRNA gene was amplified with primers 515f-m and 806r amended with Nextera tail sequences to allow subsequent barcoding (forward primer tail, TCG TCG GCA GCG TCA GAT GTG TAT AAG AGA CAG; reverse primer tail, GTC TCG TGG GCT CGG AGA TGT GTA TAA GAG ACA G). Polymerase chain reaction (PCR) was performed with the HotStarTaq Plus polymerase (Qiagen) with 5 min initial denaturation at 94°C, 35 cycles of 45s denaturation at 94°C, 60s annealing at 50°C, and 90s elongation, with a final elongation for 10 min at 72°C. Amplified libraries, including all DNA extraction blanks, PCR reagent blanks, and positive controls, were sent to the University of Minnesota Genomics Center (UMGC) for barcoding and sequencing on an Illumina Miseq platform (300 bp paired-end reads).

### 2.5 Enrichment culturing and microscopy

Select samples from the GA were enriched using liquid and solid thiosulfate media (Jones et al., 2017b) to attempt to enrich sulfur-oxidizing microorganisms, and on solid R2A media (BD Difco™, Franklin Lakes, NJ) to enrich and isolate heterotrophs. Enrichments were grown at room temperature, and liquid cultures were shaken horizontally. Colonies on solid media were plated at least twice to ensure isolation, and cultures were identified using direct PCR with primers 27F (5’-AGA GTT TGA TCC TGG CTC AG-3’) and 1492R (5’-GGT TAC CTT GTT ACG ACT T-3). Successful gene products were cleaned with the DNA Clean & Concentrator kit (Zymo Research Corp., Irvine, CA, USA), and end-sequenced at GENEWIZ (Azenta Life Sciences, Burlington, MA, USA).

For attempted cell visualization, PFA fixed samples were directly stained with 4′,6′-diamidino-2-phenylindole (DAPI) and viewed with an Olympus BX61 compound microscope with an Olympus XM10 camera running CellSens Dimensions software (Olympus, Japan).

### 2.6 Bioinformatics and statistical analyses

Raw fastq-formatted sequences were filtered and trimmed using Sickle (https://github.com/najoshi/sickle) to a minimum quality of 28 and length ≥100 bp. Residual adapters were removed from the 3’ end with cutadapt (Martin, 2011), and R1 and R2 paired reads were merged with PEAR (Zhang et al., 2014). Assembled sequences were dereplicated with VSEARCH v2.21 (Rognes et al., 2016), and clustered into operational taxonomic units (OTUs) at 97% similarity (with simultaneous chimera removal) with the UPARSE pipeline of USEARCH v11 (Edgar, 2013), using VSEARCH to map reads back to OTUs. Representative sequences for each OTU were classified with mother v1.36.1 (Schloss et al., 2009), using the Silva v132 database (Pruesse et al., 2007) and a confidence score cutoff of 50.

Statistical analyses were performed in R v4.2.1 (R Development Core Team, 2022) with the vegan v2.6-2 (Oksanen et al., 2017) and phyloseq (McMurdie, 2013) packages. First, raw counts of OTUs were converted to proportions by dividing by total number of sequences per library, and libraries with <10,000 reads were removed. As in Havlena et al. (2021), rare OTUs (<0.01% of the total) were removed, and proportional values were transformed using the angular transformation *b_ij_* = (2/π)*arcsine*[(*x_ij_*)^0.5^] to deemphasize overly abundant OTUs (McCune and Grace, 2002), where *x_ij_* is the object in the OTU matrix and *b_ij_* is transformed object. Non-metric multidimensional scaling (NMS) ordinations were calculated using the metaMDS() function with three dimensions (k=3). Hierarchical agglomerative cluster analyses were performed with Bray-Curtis dissimilarity and unweighted pair-group method using arithmetic averages (UPGMA). Q-mode clustering of samples was calculated all OTUs, and R-mode clustering of OTUs was performed with the top 50 most abundant OTUs. ANOSIM and PERMANOVA were performed with the vegan package using default parameters (Bray-Curtis distance and 999 permutations, with anosim() and adonis2() functions). Statistical significance for comparisons with 3 or more groups was assessed with a post hoc pairwise analysis using pairwise.adonis() (Martinez Arbizu, 2020). Diversity was calculated using richness and the Shannon-Wiener Index with functions rarefy() and diversity(), after subsampling the OTU matrix to 10,000 reads with function rrarefy().

Libraries were deposited in the Sequence Read Archive (SRA; http://www.ncbi.nlm.nih.gov/sra) under BioProject accession number PRJNA1005167.

## 3. RESULTS

### 3.1 Sample collection and field observations

Conditions in the cave are stable year-round, with average temperatures of 11 °C and 95% humidity (Hose et al., 2021). There is a limited air exchange between the cave and the surface, as the entrance and exit for visitors are constructed passageways with doors at each end of the tunnels. Temperatures vary up to 1 °C at tour route sites near the natural entrance, and are much more stable deeper within (Šebela et al., 2019). The GA is slightly warmer and less humid than the rest of the cave, averaging 11.4 °C and 83% humidity (G. Baker, *unpublished results*). A natural entrance pit near the north end of the Entrance Tunnel remains accessible to bats and other fauna (Great Basin National Park, 2019). Lehman Creek drainage on the surface contains springs and seasonal meltwater that infiltrates into most of Lehman Caves (Prudic & Glancy, 2009). Accumulated precipitation on Wheeler Peak, the local high point in Great Basin National Park, was 50.0 cm during our 2018 sampling trip and 109 cm during our June visit (SNOTEL historical data from USDA National Water and Climate Center, https://www.nrcs.usda.gov/wps/portal/wcc/home/). Wetter conditions resulted in more active drip water pools in areas off the main tour trail in 2019, particularly in the Cypress Swamp and Sunken Garden regions (Figure 1). Water samples collected from two perennial drip pools at the end of the GA in 2019 averaged pH 7.8, 10.7 °C, with conductivity of 356 µS cm^−1^. Eight water samples collected from the main cave averaged 7.8, 11.1 °C, and 436 µS cm^−1^ for pH, temperature, and conductivity, respectively (Supplementary Table S1). In all pools, total dissolved sulfur concentrations ranged from 0.36-3.89 mM, dissolved calcium was 1.05-3.19 mM, and dissolved silicon was 0.29-1.07 mM (Table S1). Gypsum, amorphous SiO_2_, and sepiolite (a hydrated Mg-Si mineral) were undersaturated in all water samples (Table S2).

White mineral deposits were the primary focus of sampling efforts in the GA. Morphology of the white deposits varies, where floor piles are comprised of unconsolidated powdery material, or platy and irregular, cm- and mm-sized broken fragments. *In situ* application of 5% hydrochloric acid (HCl) indicated some but not all of the white deposits contained calcium carbonate. Figure 2 shows representative field images of the samples collected, which include white crusts and loose piles lining the sides of the passage (Figure 2A). Some white residues had a soft, unstructured morphology consistent with descriptions of microcrystalline “moonmilk” commonly described in caves (Northup & Lavoie, 2001), and in some places were covered with harder, more crystalline crusts (Figure 2B). Other samples include white crusts with minor yellow coloration to their surface (Figure 2C), as well as dark brown and black loose powdery residues in limited areas (Figure 2D). Table 1 contains a summary of samples and mineral content.

**Figure 2.**
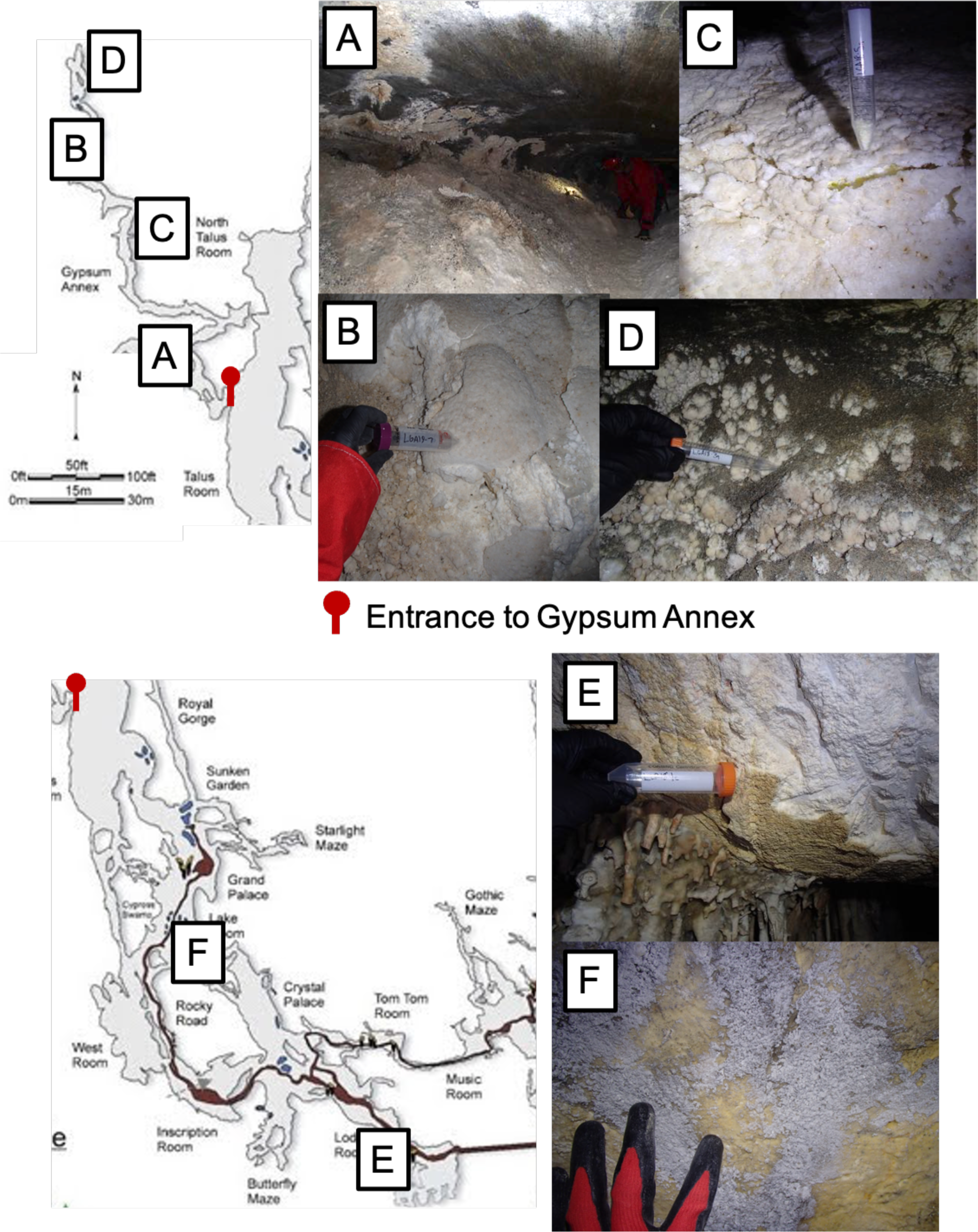
(A-D) Locations and representative photos of the substrates sampled from the Gypsum Annex. (E,F) Representative field photos and sampling locations for “main cave” samples.

We distinguish between samples collected in the GA and samples originating from other parts of the cave (“Lehman Main Cave” or LMC samples). In the main cave, we collected samples of friable material from walls and ceilings consisting of light-colored, corroded bedrock underneath a darker more resistant crust (Figure 2E). We refer to these deposits as “punk” rock, after similar corroded bedrock (Hill, 1987). Other soft substrates were sampled from ceiling locations both in GA and LMC. While these were visually similar to the rest of the surrounding rock, they could be easily poked into with a sterile metal sampling instrument, and were damp (Supplementary Figure S1A). Other samples included white/off-white, moonmilk-like coatings (example in Figure 2F), discolored ceiling material from a human-impacted area known as the “Inscription Room” (Supplementary Figure S1B), sediment from a shallow drip pool, and unconsolidated sediments.

### 3.2 Mineralogical characterization

Powder X-ray Diffraction (pXRD) showed white floor and wall deposits in the GA are composed of calcite (CaCO_3_) or gypsum (CaSO_4_•2H_2_O), and some samples have quartz (SiO_2_), dolomite (CaMg(CO_3_)_2_), and clays. Major and minor (<5%) phases for samples are summarized in Table 1. Thirteen of the 24 samples from the GA had gypsum as a major phase, and no clear spatial pattern is evident in the distribution of gypsum throughout the GA. The only other sulfate identified with pXRD was bassanite (CaSO_4_•0.5H_2_O), a sulfate hemihydrate. Anhydrite was not observed in any of the samples. Eight of the white GA deposits were predominantly calcite. Three contained high Mg-calcite as a major phase, and two contained dolomite. Seven samples from the GA, including three of the white deposits, had a minor clay component. We categorized the clays as illite or smectite group (diffraction patterns ICDD 00-058-2016 or 00-060-0320), but did not quantify them further. Three samples included hydrated magnesium silicates, either talc or sepiolite. The LMC samples were all mostly calcite.

Two samples (LGA18-5 and LGA19-13) collected from the GA were white floor crusts with several mm-sized patches of a yellow mineral. The location of these samples is indicated in Figure 3. These residues were hypothesized to be coatings of the uranyl-vanadate mineral metatyuyamunite (Ca(UO_2_)_2_(VO_4_)_2_·3H_2_O), observed in some relict SAS caves (Polyak & Mosch, 1995). SEM-EDS of a portion of the same sample mapped U and V to crystals exhibiting a platy, rhombic “booklet” habit typical of metatyuyamunite (Supplementary Figure S2) (Onac et al., 2001). We further confirmed the presence of metatyuyamunite with a 75 min pXRD scan on sample LGA19-13, as the resulting diffractogram could be manually matched to the reference pattern for metatyuyamunite (ICDD 00-043-1457).

**Figure 3.**
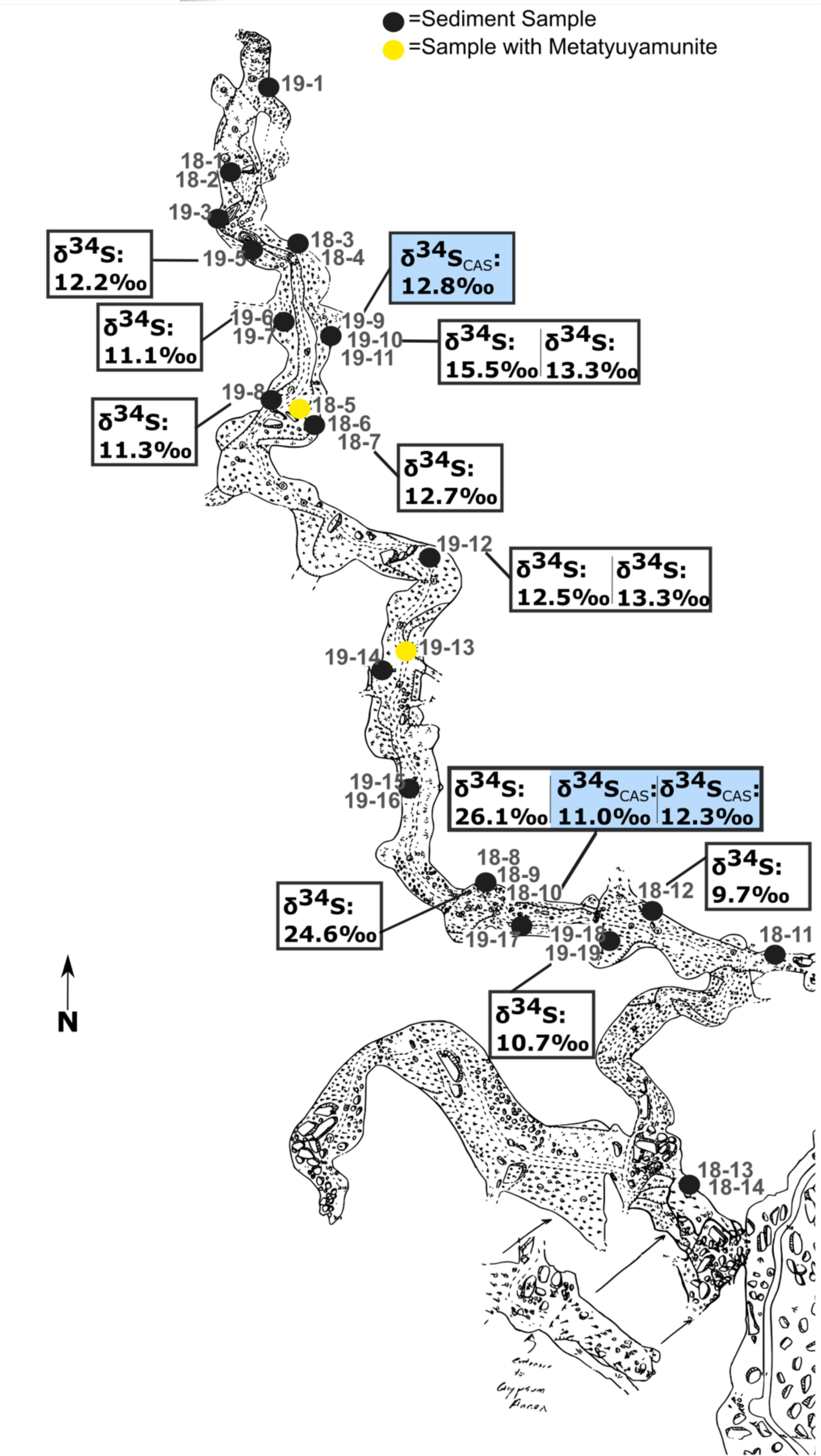
Sample locations in the Gypsum Annex (dots; hyphenated numbers are sample identifiers). Samples identified as containing metatyuyamunite are indicated in yellow, although many more have been since described throughout the passage. δ^34^S values for gypsum (δ^34^S, ±0.3‰) and carbonate-associated sulfate (δ^34^S_CAS_, ±0.4‰, blue) are shown in boxes.

Petrographic examination of two thin-sections prepared from intact crust fragments collected in the GA (LGA19-9 and LGA19-10) showed internally laminated structures not evident from the dense exterior of the hand samples. Although gypsum was identified by pXRD from the same sampling locations, the thin sections did not have evidence of gypsum. A high-relief, high order birefringent mineral consistent with calcite predominated, as well as botryoidal carbonate bands surrounded by isotropic colorless material. This colorless material displayed a texture and small-scale fractures consistent with descriptions of authigenic amorphous opal cement (Scholle & Ulmer-Scholle, 2003). Electron microprobe analysis (EMPA) with wavelength dispersive spectroscopy (WDS) confirmed the presence of Si and O (Figure 4) in the GA thin sections, consistent with disordered opal cements. EMPA also indicated heterogeneous elemental distribution within laminations, where CaCO_3_ layers were intercalated with lesser amounts of a magnesium-silicate. The only sulfur-containing mineral observed was present as a small sliver of S embedded within a layer of the mixed Mg-Si-CaCO_3_ fans.

**Figure 4.**
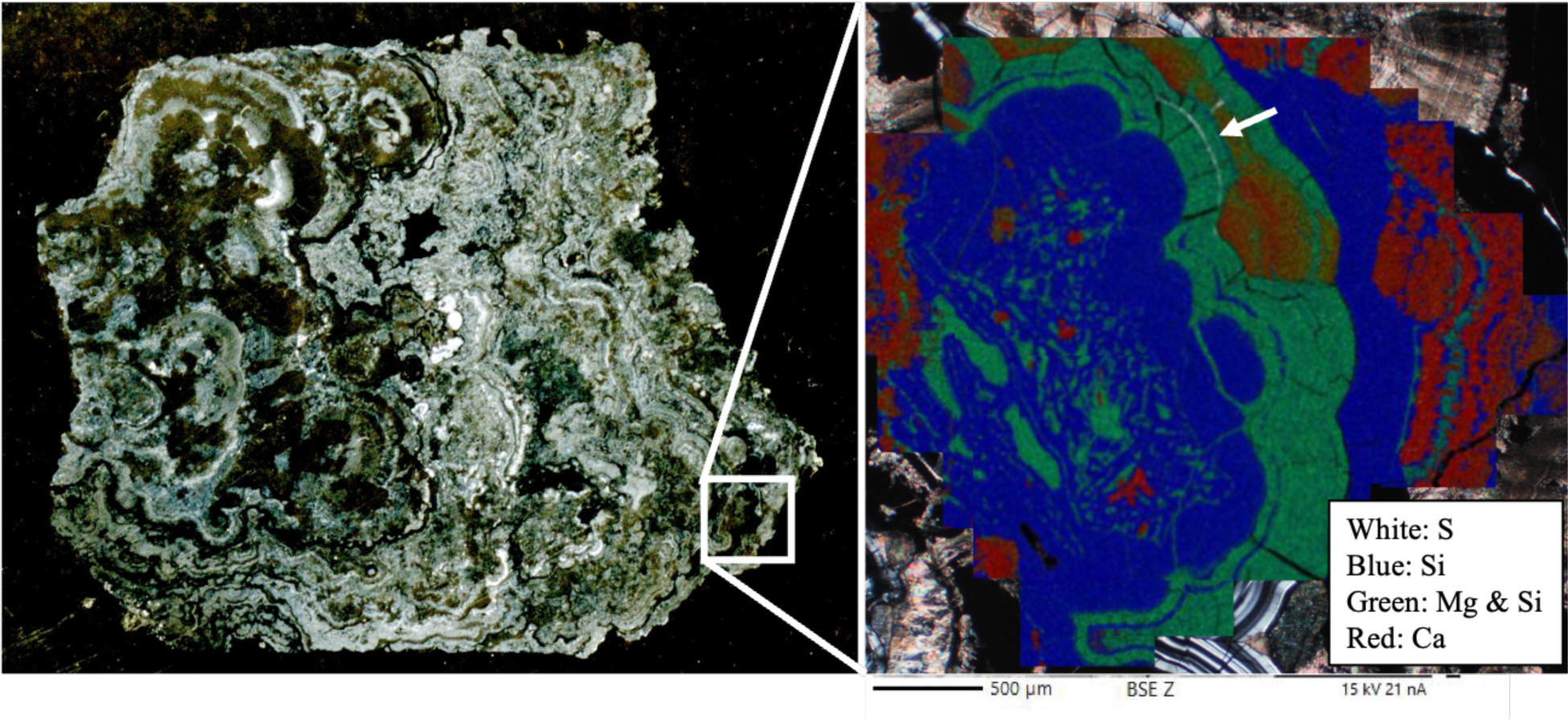
A thin section (left) from a white mineral deposit in the Gypsum Annex (sample LGA19-10). In the right panel, a photomicrograph at 5x magnification (cross-polarized) from the area indicated on the thin section are overlain with an area that was analyzed by EMPA and false colored for Si (blue), Mg (green), S (white), and carbonate (red).

### 3.3 Isotopic analysis

δ^34^S_sulfate_ values for gypsum and carbonate-associated sulfate (CAS) from the GA, summarized in Table 2, ranged from +9.7‰ to +26.1‰. Nine of the samples processed for easily soluble sulfate (ESS) and three of the samples processed for carbonate-associated sulfate (CAS) formed visible BaSO_4_ precipitates. One of the samples, LGA18-10, yielded both δ^34^S_ESS_ and δ^34^S_CAS_ values. The isotopic values for the δ^34^S_ESS_ had a wide range from +9.7‰ to +26.1‰, while CAS (δ^34^S_CAS_) values ranged from +11.0‰ to +15.6‰. The lightest δ^34^S value was +9.7‰ from sample LGA18-12, a white floor deposit collected from the south-central region of the GA, in close proximity to the second lowest value, δ^34^S +10.7 ‰, but also the two heaviest δ^34^S values (+26.1‰, and +24.6‰) (Figure 3). These heaviest values were from LGA18-8 and LGA18-10, respectively, which were collected from the surface and 10 cm apart inside the same floor breakdown deposit. However, replicate δ^34^S_CAS_ from LGA18-10 averaged +11.7, much lighter than δ^34^S_ESS_ from the same sample (14.5 ‰ lower), and more similar to ESS and CAS δ^34^S from samples collected at the north end of the passage, which ranged from +11.3‰ to +15.5‰ (Figure 3, Table 2).

**Table 2.** δ^34^S values from Gypsum Annex samples.

| Sample ID | Physical Description | $\delta^{34}\text{S}_{\text{ESS}}$<br>($\pm 0.4\text{‰}$) | $\delta^{34}\text{S}_{\text{CAS}}$<br>($\pm 0.4\text{‰}$) |
| --- | --- | --- | --- |
| LGA18-7 | Loose white floor deposit | 12.7 |  |
| LGA18-8 | Popcorn like floor breakdown pile | 24.6 |  |
| LGA18-10 | Deeper material from floor breakdown pile underneath LGA 18-8 | 26.1 | 11.0/12.3<sup>a</sup> |
| LGA18-12 | White floor deposit | 9.7 |  |
| LGA19-5 | Speleothem fragment from drip pool |  | 12.3 |
| LGA19-6 | White crust overlaying loose white floor deposit (LGA19-7) | 11.1 |  |
| LGA19-8 | Clear/white crystallization on floor breakdown | 11.3 |  |
| LGA19-9 | Powdery popcorn like fragments from wall |  | 12.8 |
| LGA19-10 | Popcorn like fragment on floor beneath LGA19-9 | 13.3/15.5<sup>a</sup> |  |
| LGA19-12 | Loose white floor deposit | 12.5/13.3<sup>a</sup> |  |
| LGA19-19 | Loose white powder on wall adjacent to LGA19-18 | 10.7 |  |

<sup>a</sup>Values from replicate samples collected at this site1041

### 3.4 Culturing and fluorescence microscopy

We identified a total of 10 morphologically distinct colonies that were isolated on solid R2A media: 8 isolates from GA samples, and 2 isolates from two samples from Lower Lodge Room (LLR) along the tour trail. R2A isolates were identified as members of the Actinobacteria and Alphaproteobacteria, including *Arthrobacter* sp., *Sphingobium* sp., *Streptomyces* sp., and *Nocardia* sp. (Table S3), indicating that there are some viable organoheterotrophic populations in the white sediments. We attempted to enrich autotrophic inorganic sulfur oxidizers from 10 GA samples, but no growth was observed on liquid or solid thiosulfate media after 3 months.

Epifluorescence microscopy of these samples was challenging. Cell counts were not performed because very few, if any, cells were visible with DAPI-staining, in part because of very low cell numbers and in part because of autofluorescent minerals and other particles from the samples.

### 3.5 Small-subunit 16S rRNA gene libraries

In initial rRNA gene libraries prepared from samples collected in 2018, over half of the GA libraries were similar to DNA extraction blanks in multivariate analyses. We therefore reevaluated DNA extraction procedures by extracting gypsum spiked with an isolate and a mock microbial community. We recovered the most DNA using the bead-beating protocol with the PowerSoil Pro DNA isolation kit (Supplementary Figure S3), and found that, surprisingly, gypsum dissolution did not improve DNA recovery. Libraries from the mock community spike in experiment prepared with the bead-beating protocol were slightly less consistent compared to some of the other protocols (Supplementary Figure S4, Supplementary Table S4); however, we selected the bead beating method given the tradeoff of recovering substantially more DNA from the low biomass samples. We then re-extracted and sequenced all samples using this protocol, and sequenced all blanks alongside sample libraries. Lehman Caves samples processed with multiple methodologies (bead beating, vortexing, and/or pre-treatment with 1x PBS for gypsum dissolution prior to extraction) that produced libraries with >10,000 sequences were included in clustering and NMS analyses.

Using this new protocol, we generated 109 16S rRNA gene amplicon libraries from samples collected during the 2018 and 2019 field expeditions, including DNA extraction blanks, PCR blanks, and positive controls. Following quality filtering and trimming, and after excluding libraries with <10,000 reads, 50 libraries had between 10,007 and 53,409 reads, with average libraries sizes 21,696±10,399. Samples with successful libraries are indicated in Table 1.

No amplification was observed in any blanks by agarose gel electrophoresis; nevertheless, four DNA extraction blanks and two PCR blanks produced libraries with more than 10,000 sequences. In two-way cluster analysis (Figure 5), most of these blank libraries occur in a cluster with two GA libraries (Cluster I in Figure 5). In this cluster, *Beijerinckiaceae* (OTU 1), *Archangiaceae* (OTU 68), *Staphylococcaceae* (OTU 13), *Chitinophagaceae* (OTU 8), and *Streptococcaceae* (OTU 17) account for 25-50% of the blanks. Some of these families are found in soil, but are also common kit contaminants (Eisenhofer et al., 2019; Salter et al., 2014), and these OTUs are not abundant in most libraries outside of Cluster I.

**Figure 5.**
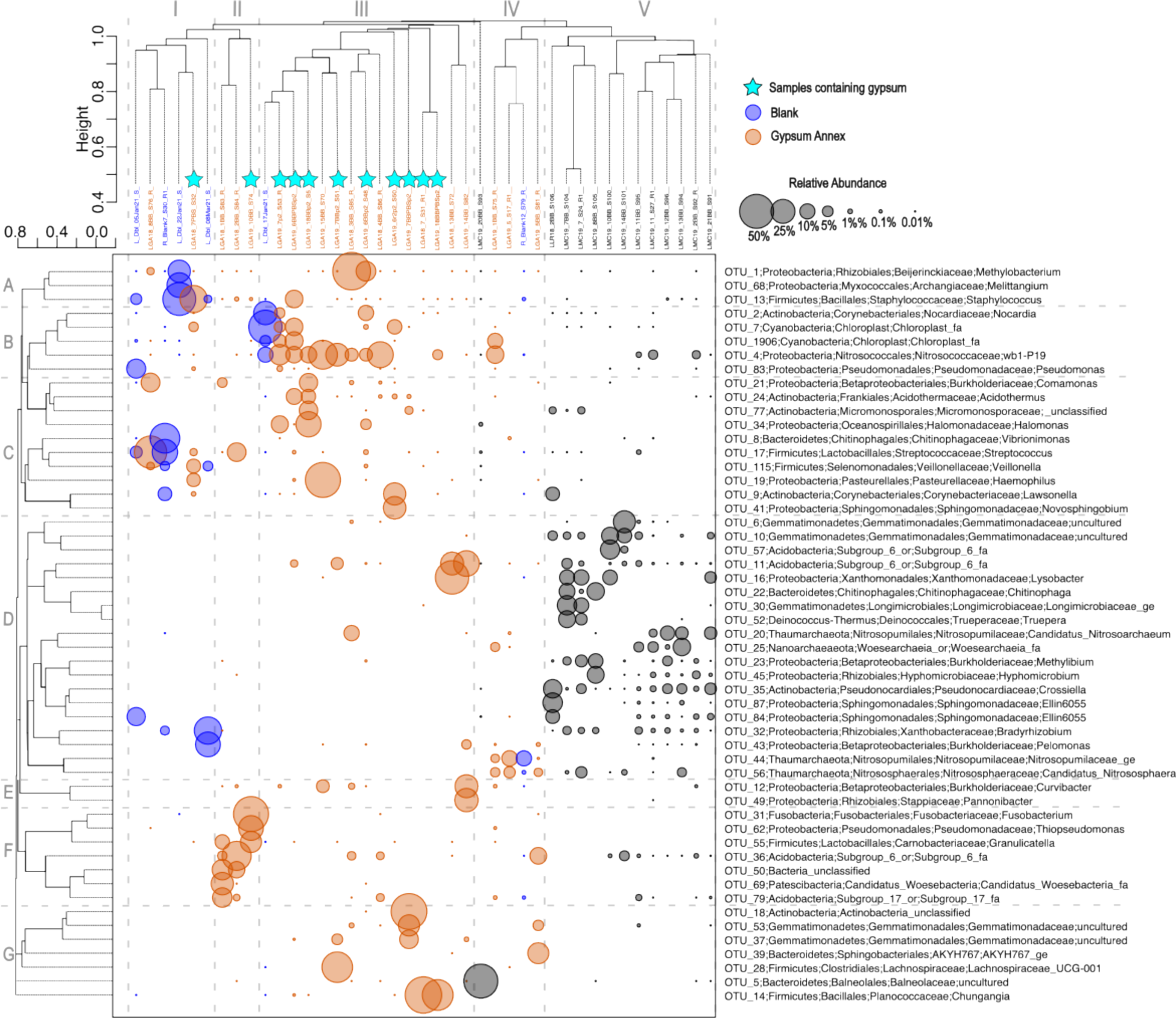
Two-way hierarchical agglomerative cluster analysis. Q-mode cluster analysis (top) was computed with all OTUs, and R-mode cluster analysis (side) was performed with only the top 50 most abundant OTUs. Samples containing gypsum are indicated with teal stars.

When blanks and the samples from Cluster 1 were removed, NMS shows that libraries from GA samples separate from libraries from the main cave along the first ordination axis (Figure 6A). GA and LMC libraries are statistically significantly different (ANOSIM, R = 0.45, p < 0.001; PERMANOVA, F = 2.8, p < 0.001). Inclusion of Cluster I libraries and blanks does not affect the overall ordination structure (Supplementary Figure S5). We also evaluated whether community composition correlates with mineralogy and sample type. In NMS ordinations, libraries from punk rock and corrosion residues separate from samples with gypsum as a major phase in ordinations, whether all samples or only samples from the GA were included (Figure 6B,C). However, these differences are not statistically significant. While ANOSIM and PERMANOVA are statistically significant (ANOSIM, R = 0.21, p = 0.03; PERMANOVA, F = 1.2, p = 0.03), post-hoc testing revealed that the only statistically significant difference is between libraries from corrosion residues and gypsum-containing samples (PERMANOVA, F = 1.6, p = 0.012; p > 0.05 for all other comparisons).

**Figure 6.**
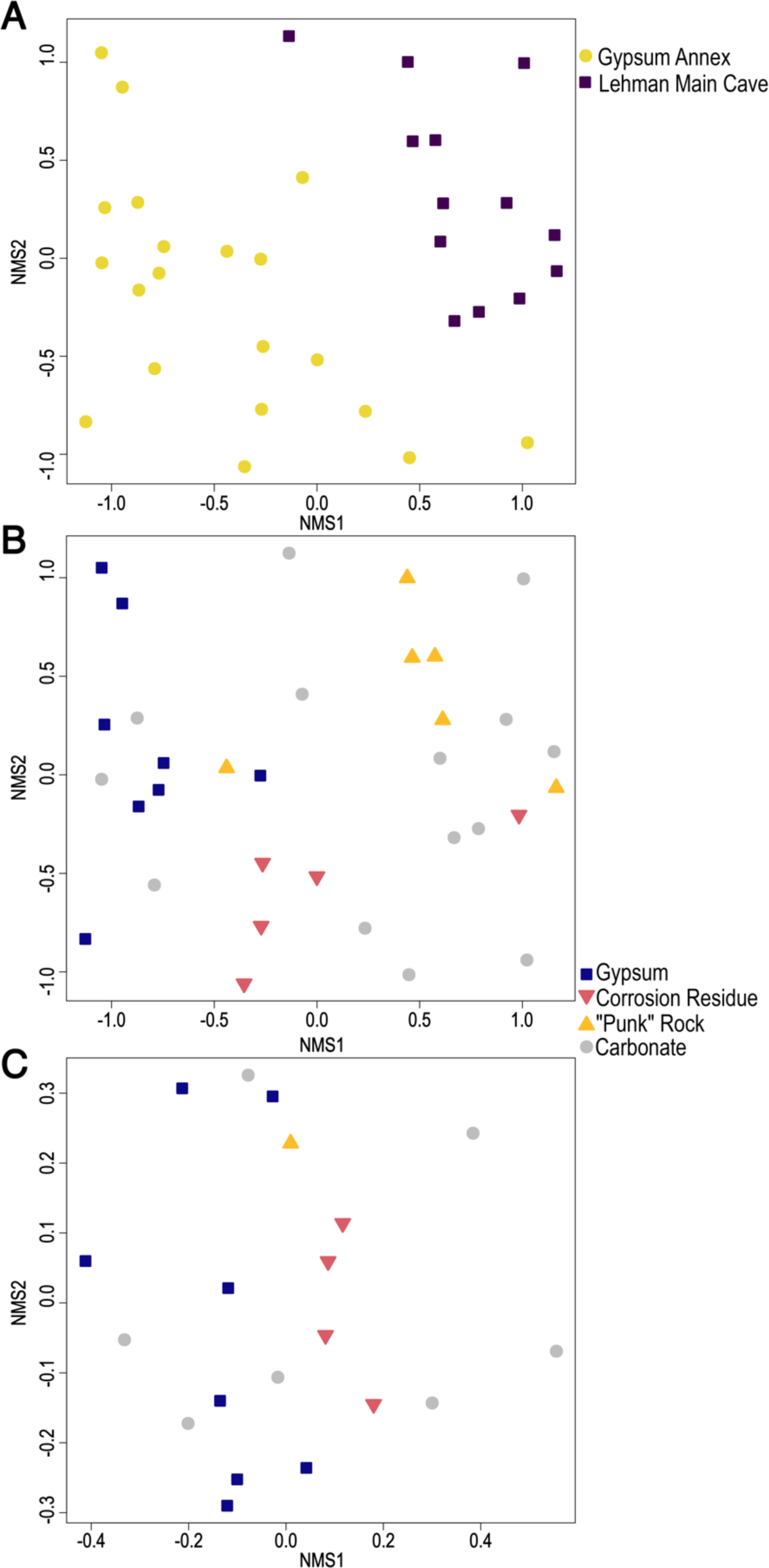
Non-metric multidimensional scaling (NMS) ordination of 16S rRNA gene libraries, with samples coded by location (panel A, Gypsum Annex versus main cave) and mineralogy (panels B and C, with all libraries in B and Gypsum Annex only in C).

Microbial taxa underlying these differences are shown in the two-way hierarchical agglomerative cluster analyses (Figure 5). Cluster III contains OTUs associated with most of the gypsum bearing deposits (indicated by stars in Figure 5). The most prevalent OTU in this cluster, OTU 4 in R-mode cluster “B”, is classified as lineage wb1-P19 in the *Nitrosococcales* (Gammaproteobacteria), and is consistently among the most abundant bacterial (≥10%) OTUs among GA samples. Clusters IV and V contain all LMC samples, where all but one fall in Cluster V. No OTUs in Cluster V are more than 25% relative abundance, and these libraries are slightly more diverse (Figure 7). The abundant taxa in these samples, in R-mode cluster D, include several bacterial OTUs from phyla Acidobacteriota, Proteobacteria, and Gemmatimonadota. Samples in Clusters IV and V both contain OTUs classified as the inorganic nitrogen-oxidizing archaeal family *Nitrosopumilaceae* (OTUs 20, 44, and 56). OTU 20 is abundant in five samples from LMC, OTU 44 occurs in samples from the GA and LMC, and OTU 56 is only in two GA samples (Figure 7).

**Figure 7.**
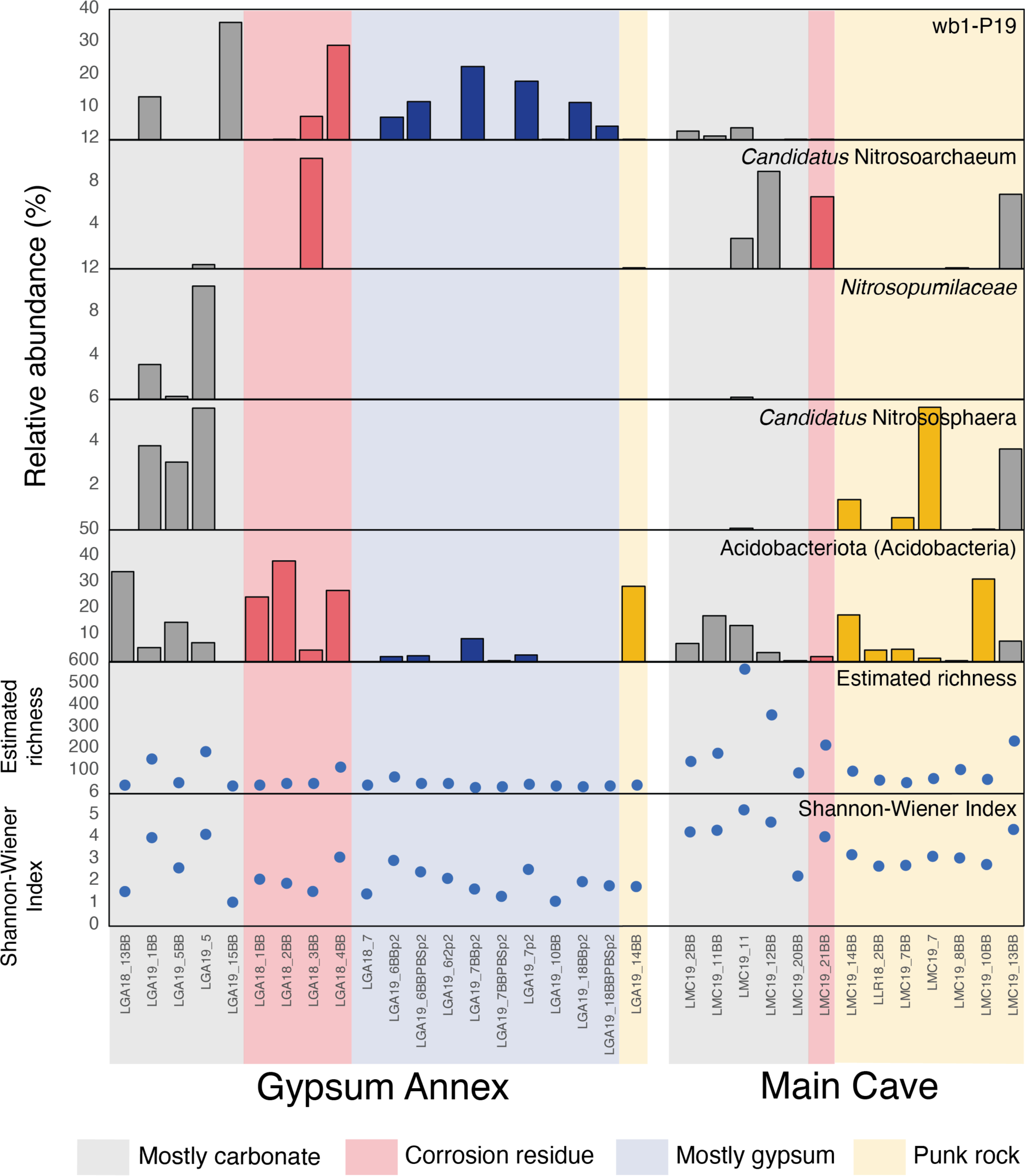
Bar chart showing the relative abundance of select taxa in cave samples by location and substrate.

Blotchy pink and orange discoloration was observed on the ceiling above the tour trail in the “Inscription Room” (Figure 1), so samples (LMC19-7 and LMC19-8) were collected to investigate if there are microorganisms that might be contributing to the discoloration. The most abundant OTUs in these samples include *Chitinophaga*, named for chitinolytic capabilities (Wieczorek et al., 2019), which is exclusively seen in these two samples of discolored ceiling substrate at 1-10% relative abundance. Other abundant OTUs in these samples were classified as genera associated with organotrophy and methylotrophy, such as *Truepera*, *Hyphomicrobium*, and *Methylibium*.

## 4. DISCUSSION

### 4.1 Origin of gypsum in Lehman Caves

White deposits lining the floors and walls of the GA are one of the most conspicuous features of the passage. However, while gypsum is present in many of the white mineral deposits within the GA, it is not ubiquitous. About 40% of the GA samples did not have detectable gypsum, and were instead predominately either calcite or high Mg-calcite. They were also associated with abundant silica, and both microcrystalline quartz as well as amorphous SiO_2_ and apparent opal cements were observed in thin section and SEM (Figure 4, Supplementary Figure S2).

The δ^34^S values from Lehman Caves gypsum are heavy compared to gypsum from known sulfidic hypogene caves. SAS gypsum from active systems such as the Frasassi Caves range from −8‰ to −24‰ (Mansor et al., 2018), +0.7‰ to −7.5‰ in Provalata Cave (Temovski et al., 2018), and 0‰ to −30‰ in Guadalupe Mountain caves (Polyak et al., 2006). In these cases, isotopically light speleogenetic gypsum reflects the initial hypogene H_2_S source. In fact, heavy δ^34^S signatures in gypsum from caves in the Nullarbor Plain (+17.1 to +21.6) were interpreted by Lipar et al. (2019) against an SAS origin as they reflected surficial sulfates. However, heavier δ^34^S can be produced by mixing processes or deep sources, such as magmatically derived sulfides, which commonly range between 0‰ to +20‰ and can confound interpretations (e.g., Onac et al., 2011, Lueth et al., 2005).

Measured δ^34^S values of ESS and CAS in the GA have a wide range of over 16‰, from +9.7‰ to +26.1‰. There also appears to be spatial differences among the ESS values, with δ^34^S_ESS_ ranging from +11.3‰ to +15.5‰ in the northern end of the passage and both of the heaviest values (+24.6 and +26.1‰) and the lightest values (+9.7 and +10.7‰) near the south-central area (Figure 3). This range suggests either multiple distinct sulfate sources or substantial fractionation during sulfur recycling. Indeed, the heaviest δ^34^S values of +26.1‰ were recovered from the ESS fraction of LGA18-10; however, δ^34^S of CAS from this and nearby samples were much lighter, +11.0‰ and +12.3‰; more similar to δ^34^S values from the northern end of the passage (Figure 3). This suggests different sulfate sources at that location during periods of carbonate and sulfate precipitation.

These “heavy” δ^34^S values are consistent with those from sulfur-bearing minerals in the greater Snake Range region. Studies have described sulfide minerals from nearby mining districts, including specimens from the Pioche District in Lincoln County, NV, about 120 km away and hosted in the same Pioche Shale and Prospect Mountain Quartzite formations that are adjacent to Lehman Caves (Vikre & Browne, 1999). δ^34^S values of these minerals fall within the +12.8‰ to +16.8‰ range seen in the GA, although none ranged as high as the two heaviest Lehman Caves samples (LGA18-8 and LGA18-10, 25.6‰ and 26.1‰, respectively). Ore depositing fluids in another eastern Nevada mineral deposit are isotopically heavier, averaging around +20‰ (Arehart et al., 1993). Another prominent sulfate source in the eastern-central Great Basin region is the Devonian bedded barites that occur mostly west of the Snake Range, and have δ^34^S isotope from 0‰ to +60‰, with most averaging around +20‰ to +40‰ (Arehart et al., 2013). While localized sulfide minerals such as pyrite can and do contribute to isotopically light gypsum in caves (Onac et al., 2011; Metzger et al., 2015), data supporting bedrock-hosted S minerals from the immediate region are not available, and further exploration of potential historical sulfide or sulfate sources around Lehman Caves is needed.

Based on mineralogical and isotopic evidence, the most likely explanation for the secondary mineral deposits in the GA is precipitation during later stage infiltration, rather than an early phase of sulfuric acid speleogenesis. While we cannot rule out that heavier δ^34^S values in the passage originated from a magmatic sulfide source, the co-occurrence of sulfates with carbonates suggests that they were not formed under the acidic conditions typical of SAS gypsum. Replacement gypsum from SAS can sometimes preserve bedrock structure (Palmer & Palmer, 2012), but that was not evident in the two samples analyzed here. Indeed, the high silica in these samples might have originated from the Prospect Mountain Quartzite, which crops out on the surface above the GA (Figure 1; Hose et al., 2021). However, a later stage origin of these secondary deposits also does not exclude a sulfuric acid origin for the cave. The hypogenic and sulfuric acid model for Lehman Caves described by Hose et al. (2021) was based on ancient morphological features, including an acid pool basin, pseudoscallops, and hollow coralloidal stalagmites. We argue that the white floor sediments in the GA were more likely subaerial precipitates formed during a period of infiltration, which could have also removed primary mineralogical evidence for sulfuric acid speleogenesis. Although the quartzite cap would have protected the GA from substantial surface infiltration, lateral infiltration may be responsible for these later-stage minerals.

### 4.2 Other secondary mineral deposits

Condensation corrosion can play an important role in speleogenesis, especially in arid caves such as Lehman Caves (Dublyansky & Dublyansky, 2000; Dublyansky et al., 2017). Traditional explanations for condensation corrosion involve CO_2_-rich airflow leading to carbonic acid etching of cave surfaces. However, in some cases, a biological role has also been suggested, such as for ferromanganese deposits (FMD), which include an oxide-rich surface and deeper “punk rock” layer (Spilde et al., 2005). In the first use of the term, Hill (1987) reported friable punk bedrock Carlsbad Caverns, and proposes that corrosion leads to crumbly, darkened cave walls and flaky residues. Black and brown loose residues within the GA (LGA18-1 and LGA18-2) are consistent with descriptions of these corrosion residues. We therefore describe similar wall deposits from Lehman Caves as punk rock, although we did not identify any Fe or Mg phases in pXRD.

We also identified yellow deposits of metayuyamunite (Ca(UO_2_)_2_(VO_4_)_2_•3-5H_2_O), or potentially tyuyamunite (Ca(UO_2_)_2_(VO_4_)_2_•5-8H_2_O), although the former is more likely given the dry conditions. Metayuyamunite has been previously identified on gypsum and dolomite in relict hypogene caves in the Guadalupe Mountains, New Mexico, USA (Polyak & Mosch, 1995), and in Caverns of Sonora, Texas, USA (Onac et al., 2001a). A definitive mechanism linking sulfidic cave processes and uranyl vanadate mineral deposition has yet to be determined. Polyak and Mosch (1995) and Polyak and Provencio (2001) proposed that U and V were liberated during sulfuric acid alteration of clays and other minerals, and metatyuyamunite precipitation occurred as surface coatings during late-stage dewatering of the cave. However, metatyuyamunite also occurs in caves and other environments that are not known to have sulfuric acid influence (Onac et al., 2001a; Onac et al., 2001b). Interestingly, botryoidal opal coatings are described as precipitating with metatyuyamunite by both Onac et al. (2001a) and Polyak and Provencio (2001).

Hydrated Mg-Si phases (talc and sepiolite) were identified in three samples by pXRD. Additionally, the Mg and Si composition of the Mg-Si phase identified by EMPA in sample LGA19-10 are consistent with sepiolite (Supplementary Figure S6). Sepiolite was not identified by pXRD of this sample, perhaps because of sample heterogeneity or poor crystallinity of the Mg-Si phase, but was identified in pXRD of LGA19-19. Sepiolite can precipitate from alkaline, Si- and Mg-rich solutions (Tosca et al., 2011; Tosca & Masterson, 2014), and has been identified as a secondary precipitate associated with carbonate phases in other caves (e.g., Martín-Pérez et al., 2021). Talc can form as a diagenetic product from sepiolite and other more hydrated and poorly crystalline Mg-Si phases (Tosca et al., 2011).

### 4.3 Mineral-associated microbial communities

Most mineral deposits had very low microbial biomass, as evidenced by our initial struggles with DNA extraction and fluorescence microscopy. After evaluating several different DNA extraction procedures, we had the best DNA recovery using the Powersoil Pro kit with bead beating, and that including a gypsum dissolution step did not noticeably improve the outcome. Overall, the mock community recovered was similar when the kit was used with bead-beating versus vortexing, with more DNA recovered when bead beating (Supplementary Figures S3 and S4).

The low biomass is probably due in part to limited input of allochthonous materials into the GA. Access to the passage is restricted except for research and surveying activities, and the sole evidence for meteoric infiltration is two ephemeral drip pools. Macrobiota are present in Lehman Caves, and include bats (typically Townsend’s big-eared bats) and troglodytic invertebrate species. Most fauna are in the main cave, but during the 2019 sampling trip, we observed a Great Basin pseudoscorpion (*Microcreagris grandis* Muchmore, endemic to Lehman Caves) in the GA, indicating that there are at least some macrobiota in the GA passage.

Some of the most abundant members of the mineral-associated microbial communities are potential ammonia oxidizing bacteria and archaea. *Nitrosococcales* lineage wb1-P19 (OTU 4), seen in over half of GA samples, was initially characterized as a possible chemolithotroph (Holmes, 2001) and has been since been described by others as having N cycling roles (Martin-Pozas et al., 2022). wb1-P19 has been identified in diverse cave types, including Weebubbie Cave in Australia (Holmes et al., 2001), a hypogene SAS cave in Italy (Jurado et al., 2020), and even volcanic caves (Weng et al., 2022). Top BLAST matches to the representative sequence of OTU 4 include several cave related sequences such as moonmilk from a limestone cave, cave biofilms, and microbial mats from lava tubes (accession numbers LT798869.1, HE602908.1, LR130706.1, OK182161.1, and KC331752.1). In addition to wb1-p19, OTUs classified as *Nitrosococcales* and three ammonia oxidizing archaea in the Thaumarchaeota (“*Candidatus* Nitrosoarchaeum,” an unclassified *Nitrosopumilaceae*, and “*Candidatus* Nitrososphera”) in GA and LMC samples further suggest that dissimilatory oxidation of inorganic nitrogen compounds may support life in diverse sediment types (Figure 7). Metagenomic analysis of these potential nitrogen-cycling organisms, or surveys of ammonia monooxygenase (*amoA*) or other genes for nitrogen compound oxidation, could improve our understanding of nitrogen cycling microorganisms in Lehman Caves and oligotrophic cave environments more generally.

We found multiple occurrences of friable bedrock and loose material that we describe as “punk rock” and other corrosion residues. These deposits included abundant OTUs classified as Acidobacteriota (formerly Acidobacteria) Subgroups 6 and 17. Subgroup 6 (OTU 36) is present in nearly all of the friable substrates sampled from both the GA and LMC. These Acidobacteriota subgroups inhabit soils world-wide, where they are implicated in degradation of soil carbon and potentially oligotrophic lifestyles (Kielak et al., 2016, de Chaves et al., 2019). Acidobacteriota make up ∼7-17% of the microbial population proposed to contribute to mineral dissolution and precipitation in cave vermiculations (Addesso et al., 2020). In Lehman Caves, Acidobacteriota could be exploiting microhabitats on mineral surfaces, or perhaps contributing more directly to carbonate degradation and formation of punk rock.

In contrast, microbial communities from sediments adjacent to the main tour trail, particularly in heavily human impacted areas such as the Inscription Room, likely reflect higher organic inputs. Microbial communities from the main cave are distinct from those from the GA (Figure 6A). Organisms in these samples include *Chitinophagaceae* and several soil associated genera in the Actinobacteria and Proteobacterial (Cluster V in Figure 5). We assume that the restricted access to the GA is at least partially responsible for these differences. Factors contributing to these differences could include elevated input of meteoric waters into the main cave, differences in speleothem texture or morphology, lint and cellular debris from tourists, legacy material from the construction of tour trails, or the presence of lighting systems. Photosynthetic microbial biofilms known as lampenflora have been previously described near artificial lights along the tour trail in Lehman Caves (Burgoyne et al., 2021). Continued protection of the GA and similar less impacted cave systems is important for preserving these environments and our ability to learn more about subsurface geomicrobiological interactions.

### 4.4 Geobiological and astrobiological significance

Gypsum and other sulfate minerals are of special interest to astrobiologists as targets for past or present life detection, particularly on Mars, which has abundant sulfate deposits (Benison & Karmanocky, 2014; Finkel et al., 2023). Caves are an attractive setting to explore the microbial ecology and biosignature preservation potential of gypsum because dark, subterranean settings are removed from photosynthetic inputs that can overwhelm biomarker detection (Johnson et al., 2020). However, our understanding of how microorganisms colonize gypsum is limited, especially in the subsurface. The gypsum-rich deposits in Lehman Caves host sparse archaeal and bacterial populations, and the low biomass of these communities might not leave a strong organic biomarker record. While we don’t know if or how active these populations are, culturing shows that at least some of these microorganisms are viable. Mineral substrate may explain some of the variability among microbial communities (Figure 6B,C), although cave location seems to be a more significant factor, which we hypothesize is related to energy resource availability. We also note that none of the microbial populations identified on either gypsum-rich or other substrates are known sulfur oxidizing or reducing chemolithotrophs, so these gypsum-associated microbial communities may not be using chemical energy from the inorganic sulfur substrate. We did identify diverse microbial populations associated with punk rock and corrosion residues, which, as other studies have argued (e.g., Northup et al., 2003; Spilde et al., 2005), may play a role in the formation of these features. Future studies should address the genomic potential and *in situ* activity of communities in Lehman Caves to further our understanding mineral substrate use and survival of cave microorganisms in this oligotrophic environment.

## Acknowledgements

Special thanks to the staff at Great Basin National Park for facilitating this research. We thank P. Scholle and D. Ulmer-Scholle for expert advice and assistance with petrographic analyses, N. Iverson, K. McNamara, and K. Hobart for their time and support with electron microscopy and XRD, and H. Aronson for providing the ICP-OES analyses. C. Dirx and other staff at the University of Minnesota Genomics Center provided insightful discussion and assistance with amplicon sequencing of challenging samples. DSJ and ZEH thank New Mexico Tech, the National Cave and Karst Research Institute (NCKRI), and NASA Exobiology Program (80NSSC20K0619) for supporting this research. ZEH thanks the Rocky Mountain Association of Geologists for supporting her with the Gary L. Babcock Memorial Scholarship. Samples were collected under National Park Service permits GRBA-2018-SCI-0012 and GRBA-2019-SCI-0010.

