## Supplementary Figures and Tables for "Origin and modern microbial ecology of secondary mineral deposits in Lehman Caves, Great Basin National Park, NV, USA"

**Table S1.** Geochemical data from cave pools

| Sample | pH | Temp ($^{o}C$) | Conductivity (µS/cm) | Al (mM) | Ca (mM) | Fe (mM) | K (mM) | Mg (mM) | Mn (mM) | Na (mM) | P (mM) | S (mM) | Si (mM) | Sr (mM) |
|---|---|---|---|---|---|---|---|---|---|---|---|---|---|---|
| Gypsum Annex | | | | | | | | | | | | | | |
| LGA19-2b | 7.74 | 10.3 | 350 | 0.002 | 1.981 | nd | 0.034 | nd | nd | 0.554 | nd | 0.611 | 0.285 | nd |
| LGA19-4b | 7.77 | 10.4 | 342 | nd | 2.144 | nd | 0.041 | nd | nd | 0.644 | 0.001 | 0.704 | 0.365 | nd |
| LGA19-20b | 7.75 | 11.3 | 376 | nd | 1.19 | nd | 0.05 | nd | nd | 0.583 | nd | 0.691 | 0.572 | nd |
| Main Cave | | | | | | | | | | | | | | |
| LMC19-1a | 7.84 | 11.3 | 370 | nd | 1.521 | nd | 0.081 | 0.028 | nd | 0.773 | nd | 0.862 | 0.787 | nd |
| LMC19-3b | 7.78 | 11 | 274 | nd | 1.049 | nd | 0.031 | nd | nd | 0.26 | nd | 0.359 | 0.458 | nd |
| LMC19-4b | 7.75 | 11.3 | 810 | nd | 1.621 | nd | 0.082 | 0.077 | nd | 2.913 | 0.001 | 3.887 | 1.067 | nd |
| LMC19-5b | 7.9 | 11 | 340 | nd | 1.33 | nd | 0.043 | nd | nd | 0.261 | nd | 0.499 | 0.546 | nd |
| LMC19-15 | 7.81 | 11.1 | 442 | nd | 1.444 | nd | 0.117 | nd | nd | 0.807 | 0.001 | 0.789 | 0.572 | nd |
| LMC19-17 | 7.77 | 11.2 | 438 | nd | 1.399 | nd | 0.067 | 0.003 | nd | 0.577 | 0.001 | 0.799 | 0.638 | nd |
| LMC19-18 | 7.89 | 11.2 | 406 | 0.003 | 3.185 | nd | 0.11 | 0.026 | nd | 0.832 | 0.002 | 0.969 | 0.698 | nd |
| LMC19-19 | 7.78 | 10.6 | 407 | 0.002 | 2.089 | nd | 0.028 | nd | nd | 0.24 | nd | 0.44 | 0.425 | nd |

nd = not detected

**Table S2.** Saturation indices for select mineral phases in cave pools, based on the data presented in Table S1

| Sample | Gypsum | Anhydrite | $SiO_2$(am) | Quartz | Chalcedony | Talc | Sepiolite | Hydroxyapatite |
|---|---|---|---|---|---|---|---|---|
| | $CaSO_4{\bullet}2H_2O$ | $CaSO_4$ | $SiO_2$ | $SiO_2$ | $SiO_2$ | $Mg_3Si_4O_{10}(OH)_2$ | $Mg_2Si_3O_{7.5}OH{\bullet}3H_2O$ | $Ca_5(PO_4)_3OH$ |
| Gypsum Annex | | | | | | | | |
| LGA19-2b | -1.69 | -2.16 | -0.71 | 0.66 | 0.18 | - | - | - |
| LGA19-4b | -1.61 | -2.08 | -0.6 | 0.77 | 0.29 | - | - | - |
| LGA19-20b | -1.8 | -2.26 | -0.41 | 0.95 | 0.47 | - | - | - |
| Main Cave | | | | | | | | |
| LMC19-1a | -1.64 | -2.1 | -0.28 | 1.09 | 0.61 | -2.56 | -3.53 | - |
| LMC19-3b | -2.1 | -2.57 | -0.51 | 0.86 | 0.38 | - | - | - |
| LMC19-4b | -1.11 | -1.57 | -0.14 | 1.22 | 0.75 | -1.59 | -2.83 | -1.28 |
| LMC19-5b | -1.89 | -2.35 | -0.43 | 0.93 | 0.46 | - | - | |
| LMC19-15 | -1.69 | -2.15 | -0.41 | 0.95 | 0.48 | - | - | -0.62 |
| LMC19-17 | -1.69 | -2.15 | -0.37 | 1 | 0.52 | -6.25 | -6.01 | -0.82 |
| LMC19-18 | -1.38 | -1.84 | -0.33 | 1.04 | 0.56 | -2.69 | -3.62 | 1.8 |
| LMC19-19 | -1.81 | -2.28 | -0.54 | 0.83 | 0.35 | - | - | - |

**Table S3.** Summary of 16S rRNA gene sequences from R2A isolates

| Location | Cave sample | Isolate ID | Inferred taxonomy | Top BLAST hit | Accession | Percent identity |
|---|---|---|---|---|---|---|
| Gypsum Annex | LGA18-1 | LGR-11 | Streptomyces sp. | Streptomyces cirratus strain QT193-6 | MT083993 | 99.85 |
| Gypsum Annex | LGA18-4 | LGR-18 | Nocardia sp. | Nocardia sp. strain K78 | MT422810.1 | 99.57 |
| Gypsum Annex | LGA18-5 | LGR-20 | Actinomycetia sp. | Actinomycetia bacterium strain Qhu-G193 | OP881727.1 | 99.93 |
| Gypsum Annex | LGA18-11 | LGR-21 | Streptomyces sp. | Streptomyces sp. TFS59-23 | HM001271.1 | 98.78 |
| Gypsum Annex | LGA18-7 | LGR-7 | Arthrobacter sp. | Arthrobacter sp. strain MJS-AB-C17 | MG694494.1 | 98.36 |
| Main Cave | LLR18-1 | LLR-3 | Sphingomonas sp. | Uncultured bacterium clone 33MIC093* | JF341241.1 | 98.09 |
| Main Cave | LLR18-2 | LLR-5 | Streptomyces sp. | Streptomyces sp. strain XG100 | OQ286339.1 | 98.15 |
| Gypsum Annex | LGA18-1 | LGR-2 | Rhizorhapis sp. | Rhizorhapis suberifaciens strain JCM 8524 | LC379084.1 | 96.23 |
| Gypsum Annex | LGA18-3 | LGR-12 | Nocardia sp. | Nocardia fluminea isolate 0911ARD13M2 | LN774198.1 | 97.66 |
| Gypsum Annex | LGA18-4 | LGR-9 | Streptomyces sp. | Streptomyces cirratus strain QT-155 | MT081095.1 | 98.15 |

*Top BLAST hit to a named organism is Sphingomonas sp. (MZ675813.1), 95.62% identity

**Table S4.** Mock community recovery from gypsum spike-in experiments

| Genus | Expected mock community composition | Vortexing (5, 10, 15 min intervals) | Vortexing 10 min | Bead beating (20, 40, 60 s intervals) | Bead beating, 60 s | Mock community only (no gypsum) |
|---|---|---|---|---|---|---|
| *Bacillus* | 17.4 | 16.1 (±2.2) | 14.2 (±0.7) | 10.3 (±0.7) | 9.6 (±0.8) | 10.8 (±0.3) |
| *Enterococcus* | 9.9 | 7.5 (±0.1) | 7.1 (±1.7) | 8.9 (±0.2) | 9.9 (±0.2) | 10.2 (±0.4) |
| *Escherichia-Shigella* | 10.1 | 11.4 (±1.5) | 12.9 (±1.4) | 14.8 (±0.4) | 15.5 (±0.6) | 15.7 (±0.6) |
| *Lactobacillus* | 18.4 | 21.5 (±1.3) | 22.5 (±1.7) | 21.2 (±1.0) | 22.3 (±0.4) | 19.5 (±0.3) |
| *Listeria* | 14.1 | 9.6 (±1.7) | 8.2 (±1.5) | 5.0 (±0.4) | 3.8 (±0.9) | 4.5 (±0.1) |
| *Pseudomonas* | 4.2 | 8.4 (±1.3) | 9.3 (±1.4) | 12.0 (±0.2) | 13.1 (±0.7) | 12.2 (±0.7) |
| *Salmonella* | 10.4 | 11.1 (±1.5) | 12.9 (±1.8) | 15.2 (±0.8) | 15.1 (±1.1) | 15.7 (±1.0) |
| *Staphylococcus* | 15.5 | 14.1 (±1.5) | 9.8 (±2.9) | 9.8 (±0.5) | 8.4 (±1.2) | 11.2 (±0.7) |

All extractions performed using the PowerSoil Pro kit. Numbers represent the average and standard deviation of triplicate DNA extractions

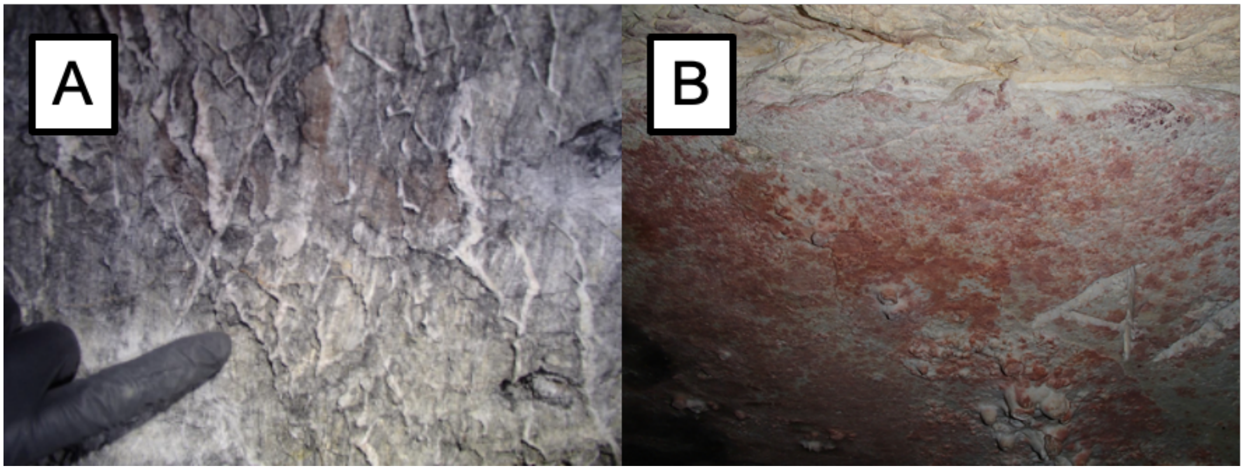

**Supplementary Figure S1:** A: Soft ceiling from sample location LGA19-14, white calcite veins were hard compared to friable gray dolomite wall rock. B: Pink discoloration and etchings of friable ceiling in Inscription Room, sample LMC19-7.

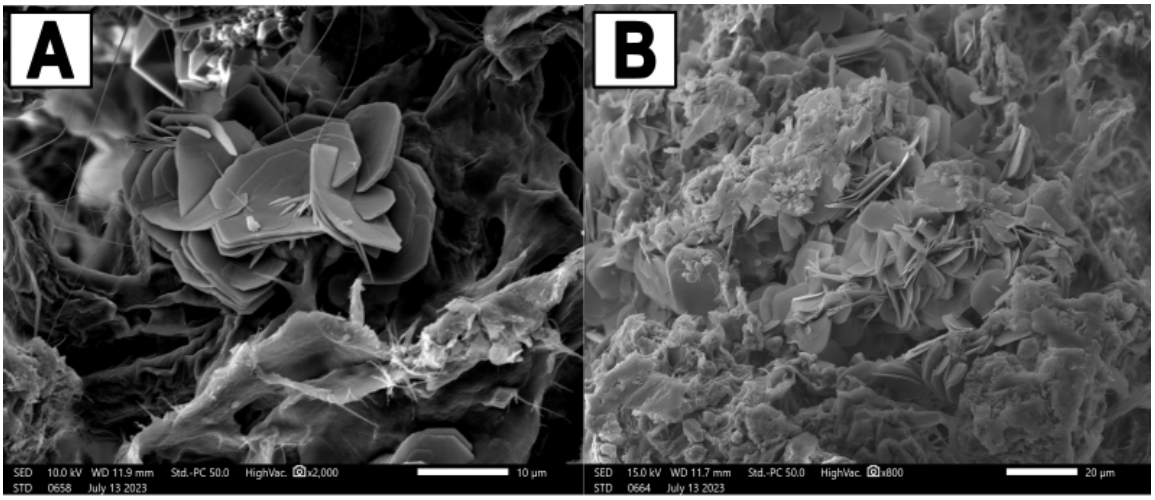


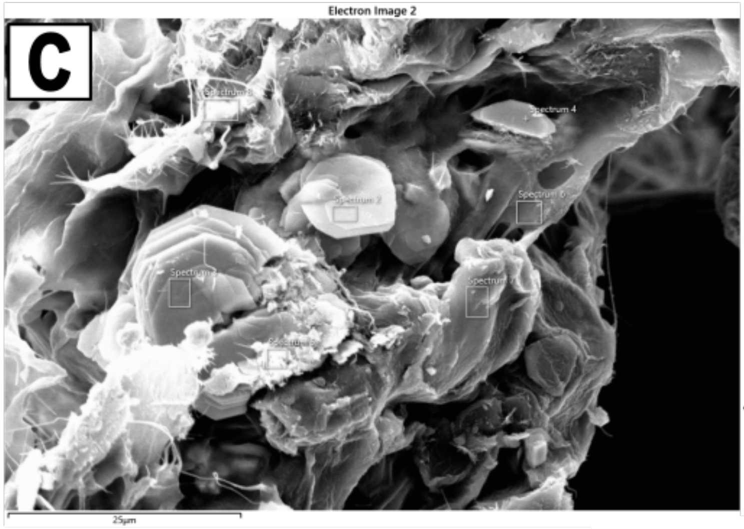


| | Spectrum2 | Spectrum3 | Spectrum4 | Spectrum5 | Spectrum6 | Spectrum7 | Spectrum8 | Spectrum9 | Spectrum10 | Spectrum11 |
|---|---|---|---|---|---|---|---|---|---|---|
| **C** | 21 | 15 | 16 | 15 | 12 | 9.5 | 12 | 23 | | 14 |
| **O** | 58 | 69 | 68 | 67 | 63 | 63 | 63 | 61 | 61 | 68 |
| **Mg** | 3.1 | | 1.8 | 3.6 | 9.6 | 11 | 11 | | 16 | |
| **Si** | 3.3 | | 2.5 | 4.4 | 15 | 16 | 14 | | 23 | |
| **K** | | | | | | | | | | 2.3 |
| **Ca** | 2.8 | 2.9 | 2.3 | 2.1 | 0.5 | 0.7 | 0.3 | 16 | 0.7 | 2.1 |
| **V** | 5.7 | 6 | 4.5 | 3.8 | | | | | | 6.3 |
| **U** | 6.2 | 7.1 | 5.3 | 4.4 | 0.2 | | | | | 7.3 |
| Total | 100 | 100 | 100 | 100 | 100 | 100 | 100 | 100 | 100 | 100 |

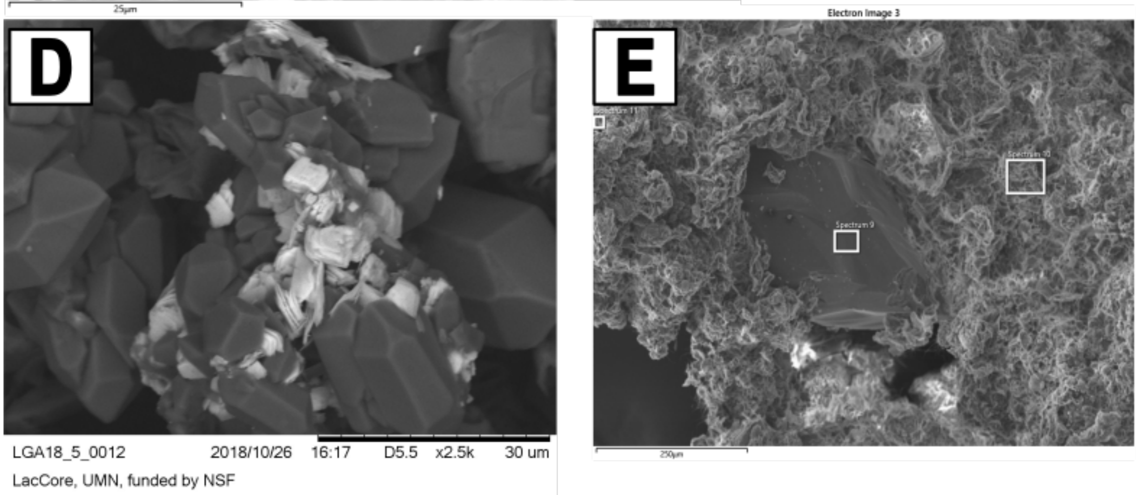


**Supplementary Figure S2:** SEM images of the yellow crust on white floor deposit sample LGA19-13 from the Gypsum Annex (images A, B, C, and E, secondary electron imaging). EDS spectra correspond to the white squares in images C and E, and show the presence of uranium and vanadium associated with the platy minerals. D, Backscattered electron image of sample LGA18-5 taken with a TM-1000 SEM.

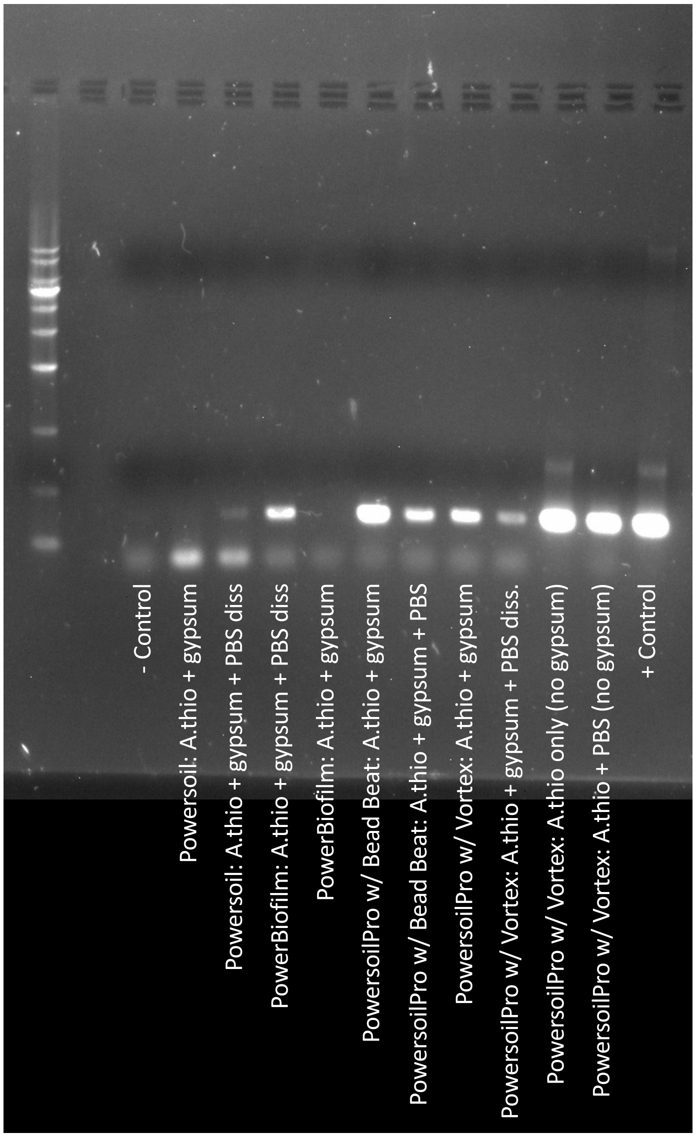


**Supplementary Figure S3:** PCR products from DNA extracts of gypsum spiked with *Acidithiobacillus thiooxidans* cells (35 PCR cycles with V4 primers). DNA was extracted with the DNeasy Powersoil kit (now discontinued), the DNeasy PowerBiofilm kit, and the DNeasy Powersoil Pro kit, the latter extracted using a Vortex-Genie 2 (for 5, 10, and 15 min at maximum speed) or a Biospec Products Mini Beadbeater (for 20, 40, and 60 s at 2500 RPMs). "PBS diss" refers to experiments in which gypsum was first dissolved by horizontally shaking the sample in 1× phosphate buffered saline (PBS), pH 2. The $2^{nd}$ and $3^{rd}$ bands from the right were extractions of an equivalent amount of *Acidithiobacillus* only, with no gypsum.

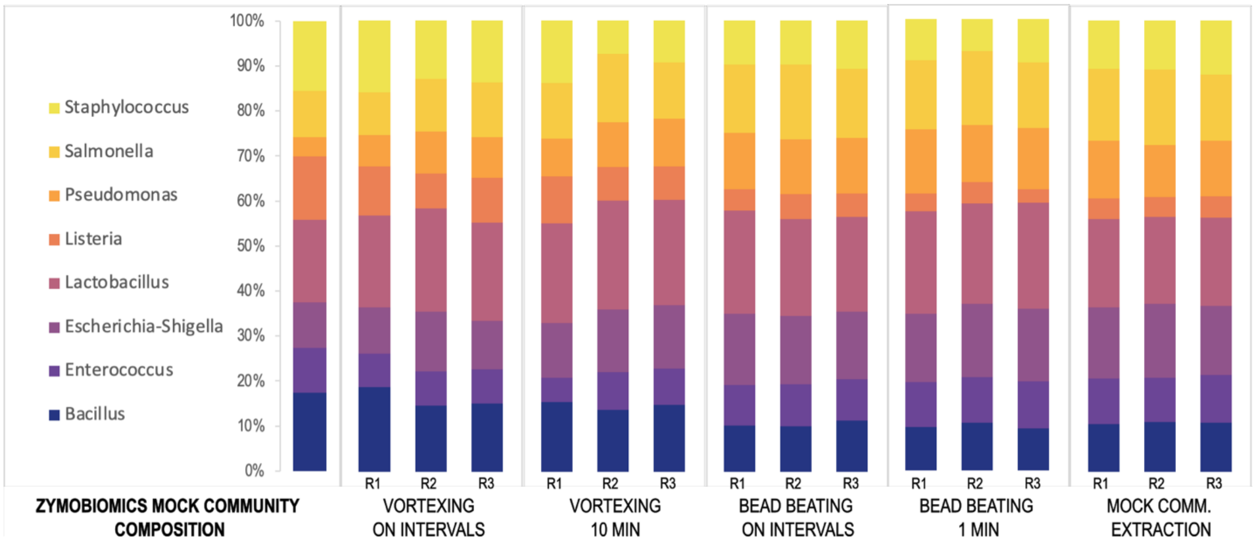


**Supplementary Figure S4.** Comparison of mock community recovery in 16S rRNA gene libraries from gypsum spike-in experiments using whole cells from the Zymobiomics mock community standard. DNA was extracted using the DNeasy PowerSoil Pro kit with either a Vortex-Genie 2 (for either 10 minutes, or 5, 10, and 15 min) or a Biospec Products Mini Beadbeater (for either 60 s, or 20, 40, and 60 s), each in triplicate. The right columns are from extraction of the same amount of mock community alone. Bars correspond to values in Table S4.

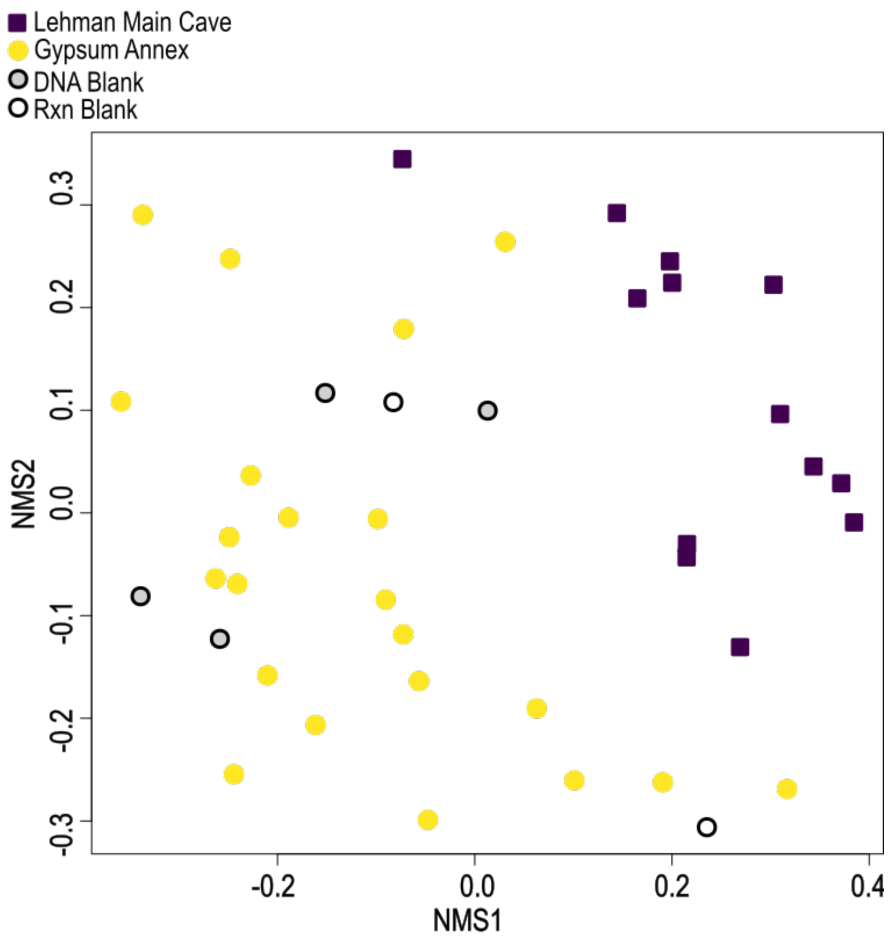


**Supplementary Figure S5:** Non-metric multidimensional scaling (NMS) ordination of 16S rRNA gene libraries, similar to Figure S7a in the main text but with DNA and PCR blanks included.

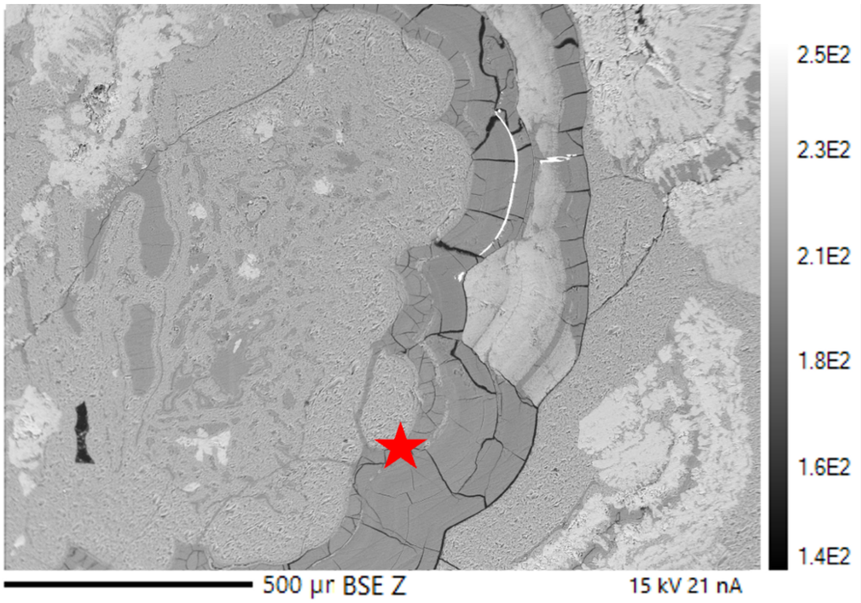


| Atomic% | | Oxide wt. % | | | |
|---|---|---|---|---|---|
| | LGA10 WDS-quant. point | | LGA10 WDS-quant. point | Martín-Pérez et al., 2021[1] | |
| Na | 0.19 | Na2O | 0.24 | | |
| F | 1.28 | F | 1 | | |
| Si | 22.44 | **SiO2** | 55.51 | 54.5 | 53.8 |
| Al | 0 | Al2O3 | 0 | | |
| Mg | 15.37 | **MgO** | 25.51 | 24.1 | 26.9 |
| P | -0.01 | P2O5 | -0.02 | | |
| S | 0.01 | SO2 | 0.02 | | |
| Cl | 0 | Cl | 0 | | |
| K | 0.02 | K2O | 0.03 | | |
| Ca | 0.18 | CaO | 0.43 | 0.8 | 0.5 |
| Ti | 0 | TiO2 | -0.01 | | |
| Mn | 0 | MnO | 0.01 | 0 | 0 |
| Fe | -0.01 | FeO | -0.02 | 0 | 0 |
| O | 60.53 | | | | |
| Total | 100 | Analytical Total | 82.75 | 79 | 81 |

[1]Martín-Pérez A, La Iglesia Á, Almendros G, González-Pérez JA, Alonso-Zarza AM (2021) Precipitation of kerolite and sepiolite associated with Mg-rich carbonates in a cave environment. *Sedimentary Geology* **411**, 105793.

**Supplementary Figure S6:** Backscattered electron (BSE) image of LGA19-10 showing Mg-Si rich phases. The wavelength dispersive X-ray spectroscopy (WDS) data shows $SiO_2$ to MgO ratios characteristic of sepiolite (at point highlighted by red star). Low analytical totals correspond to the unanalyzable water component, which is approximately 18 wt. %. $SiO_2$ and MgO data is compared to two oxide wt. % data points from Martín-Pérez et al. (2021), which identified sepiolite and kerolite associated with cave carbonates.
